# Next-level high-precision spatial omics enabled through parallel image acquisition and spatial similarity networking

**DOI:** 10.1101/2025.03.07.642022

**Authors:** Varun V. Sharma, Gabor Toth, Robert Martinis, Cathrin E. Hansen, Gijs Kooij, Ingela Lanekoff

**Affiliations:** Department of Chemistry - BMC, Uppsala University, Uppsala, Sweden; Center of Excellence for the Chemical Mechanisms of Life, Uppsala University, Sweden; Amsterdam UMC, Location VU Medical Center, Department of Molecular Cell Biology and Immunology, De Boelelaan 1117, Amsterdam, The Netherlands; Amsterdam Neuroscience, Amsterdam UMC, Amsterdam, The Netherlands; MS Center Amsterdam, Amsterdam UMC, Location VU Medical Center, Amsterdam, The Netherlands; Amsterdam Institute for Immunology and Infectious Diseases, Amsterdam UMC, Amsterdam, The Netherlands

## Abstract

Unambiguous molecular annotations are essential to discern complex local biochemical processes in spatial biology. Here we present a scalable and broadly applicable platform for tandem mass spectrometry imaging (MS^2^I) that overcomes current limitations in annotation with MSI by integrating Parallel Image Acquisition (PIA) with a novel open-access computational framework, Spatial Similarity Networking (SSN). The PIA employs parallelized acquisition of untargeted MSI and targeted MS^2^I data using multiple inclusion lists to ensure spatially consistent and structure-resolved imaging of hundreds of molecular species in a single experiment. For molecular annotation, we have developed the SSN that complements PIA by leveraging spatial correlations among product ions through a graph-based analysis framework to enable confident molecular annotation even within highly complex MS^2^I datasets. Using this integrated approach, we successfully resolved and annotated 134 phospholipid isomers and isobars from mouse brain tissue and suggest confidence levels for annotation for the MSI community. Furthermore, we applied our platform to interrogate cholesterol metabolism in human multiple sclerosis brain tissue, achieving annotation of six novel brain-related oxysterols and revealing spatially correlated oxidation pathways linked to lesion severity. Together, PIA and SSN establish a new framework for large-scale, structure-specific mass spectrometry imaging, with broad implications for spatial metabolomics, lipidomics, and chemical pathology beyond current capabilities.

## Introduction

The structures of small molecules that regulate cellular processes are highly specific, with isomeric forms often producing distinct biological effects.^1^ Structural isomers of metabolites and lipids can alter the function of cells and cellular regions by regulating key physiological processes, such as selectively interacting with enzymes or receptors, and modulating signalling pathways.^2^ Accurately annotating molecules is essential, and clear standards for molecular annotation have been adopted by the metabolomics community for bulk samples.^3–5^ Yet, annotating and spatially localizing isomers within biological tissue remains a central challenge.^6^

Mass spectrometry imaging (MSI) has emerged as a cornerstone for localizing molecules in situ by enabling highly multiplexed and label-free spatial mapping of metabolites, lipids, and other biomolecules.^7–9^ The capabilities of MSI have transformed investigations of tissue biochemistry, cellular heterogeneity, and pathological mechanisms.^10,11^ For example, MSI has revealed localized metabolic reprogramming as a key driver of cancer progression,^12^ spatially resolved metabolic interplay at the host-microbe interface,^13^ and drug distributions in pharmacological studies.^14^ These spatially resolved molecular insights have uncovered previously unrecognized chemical heterogeneity and critical molecular microenvironments that influence disease progression, metabolic regulation, and therapeutic insights unattainable through bulk analysis.

Despite the advances, a key limitation of MSI is its inability to distinguish isomeric and isobaric species, even when augmented by high mass resolution or ion mobility.^15^ Consequently, confidence in molecular annotation at isomeric level remains low in MSI workflows.^6,16^ The combination of MSI with tandem mass spectrometry (MS^2^) in selected regions adds valuable structural information and is achieved by untargeted strategies such as data-dependent and dataset-dependent acquisition workflows.^17,18^ However, untargeted strategies do not consistently fragment precursor ions across the tissue, hiding the dynamic distribution of isomers.^19,20^ This calls for consistent spatially resolved isomer imaging that can be achieved with tandem mass spectrometry imaging (MS^2^I).^21^

In MS^2^I, each pixel of the ion image contains molecular fragmentation data for annotation of isomers and isobars, and therefore, product ion images can be generated and visualized. However, in addition to the need for consistent precursor ion selection across pixels, the widespread adoption of MS^2^I is hindered by trade-offs between spatial resolution, throughput, and molecular coverage.^22–26^ For example, despite enabling MS^2^I of 92 isolation windows using a target MS^2^ list, the sequential MS^1^ and MS^2^ scans compromise throughput and spatial resolution.^23^ This can be improved by faster mass analysers with low mass resolution,^22,24,27^ although annotation based solely on low mass resolution can challenge annotation due to the wide isolation window often containing multiple precursors.

Here we present a scalable MS^2^I platform that advances MSI by integrating Parallel Image Acquisition (PIA) with the computational framework Spatial Similarity Network (SSN). PIA enables multiplexed parallel acquisition of high-mass resolution untargeted MSI and scalable targeted MS^2^I data, allowing spatially consistent, structure-resolved imaging of hundreds of molecular species with conserved pixel size and experimental time. SSN enhances molecular annotation by exploiting the distribution of product ions, computing their spatial correlations to identify isomer annotations and spatial distributions. We utilize our novel PIA SSN workflow to spatially map isomeric sterols and phospholipids in human and mouse brain, respectively. Overall, PIA and SSN together establish a framework for large-scale, structure-specific mass spectrometry imaging, advancing spatial metabolomics, lipidomics, and chemical pathology.

## Results and Discussion

### Scalable Structure-Resolved Imaging via Parallel Image Acquisition

The ambient MSI technique pneumatically assisted nanospray desorption electrospray ionization (PA nano-DESI) enables the continuous generation of high ion flux, providing exceptional sensitivity for MSI of low-abundant metabolites, such as prostaglandins and steroids, which are generally difficult to ionize (Fig. 1a).^28–30^ When combined with a hybrid mass spectrometer, PA nano-DESI allows for multistage MS^n^I, allowing for isomer differentiation and imaging in MS^2^I, MS^3^I, and MS^4^I through the sequential acquisition of individual scans.^20,22,30–32^ Integrating PA nano-DESI with a tribrid mass spectrometer, comprising a quadrupole mass filter, an Orbitrap (FT), and an ion trap (IT) mass analyser, enables parallel data acquisition. Crucially, in tribrid instruments, the Orbitrap and ion trap analyzers can operate independently, allowing MS and MS^2^ spectra to be acquired simultaneously without temporal interference.^26^ The parallel acquisition strategy enables repeated fragmentation of each precursor at every pixel, conserving pixel size while minimizing ion wastage (Fig. 1b). Specifically, a typical 1024 ms FT scan (*m/Δm* = 500,000 at *m/z* 200) with an ion accumulation time of 5 ms, >99.5% of the ions entering the mass spectrometer remain unutilized. Our approach overcomes this inefficiency, where up to 27 MS^2^ spectra, containing over 100 unique precursors, can be acquired in the IT simultaneously with one FT scan (Fig. 1c-e and Supplementary Fig. 1a). This results in a substantial increase in structural information per pixel without extending acquisition time or compromising pixel size. Moreover, our approach enables extensive molecular coverage by looping through multiple inclusion lists during acquisition (Fig. 1c). For example, cycling through four inclusion lists, each targeting 27 narrow precursor isolation windows (0.7 Da), enables the acquisition of MS^2^ data from 108 distinct isolation windows in a single imaging experiment. While looping through *N* inclusion lists nominally increases the x-dimension pixel size for ITMS^2^ by a factor of *N*, the method enables simultaneous visualisation of thousands of product ion images in one experiment (Fig. 1c and Supplementary Fig. 1b-c). We introduce this highly multiplexed workflow as parallel image acquisition (PIA). Overall, PIA enables the combination of untargeted full-scan MSI in parallel with high-molecular-coverage targeted MS^2^I for precise spatial localization of product ions in a single, scalable experiment.

**Fig 1.**
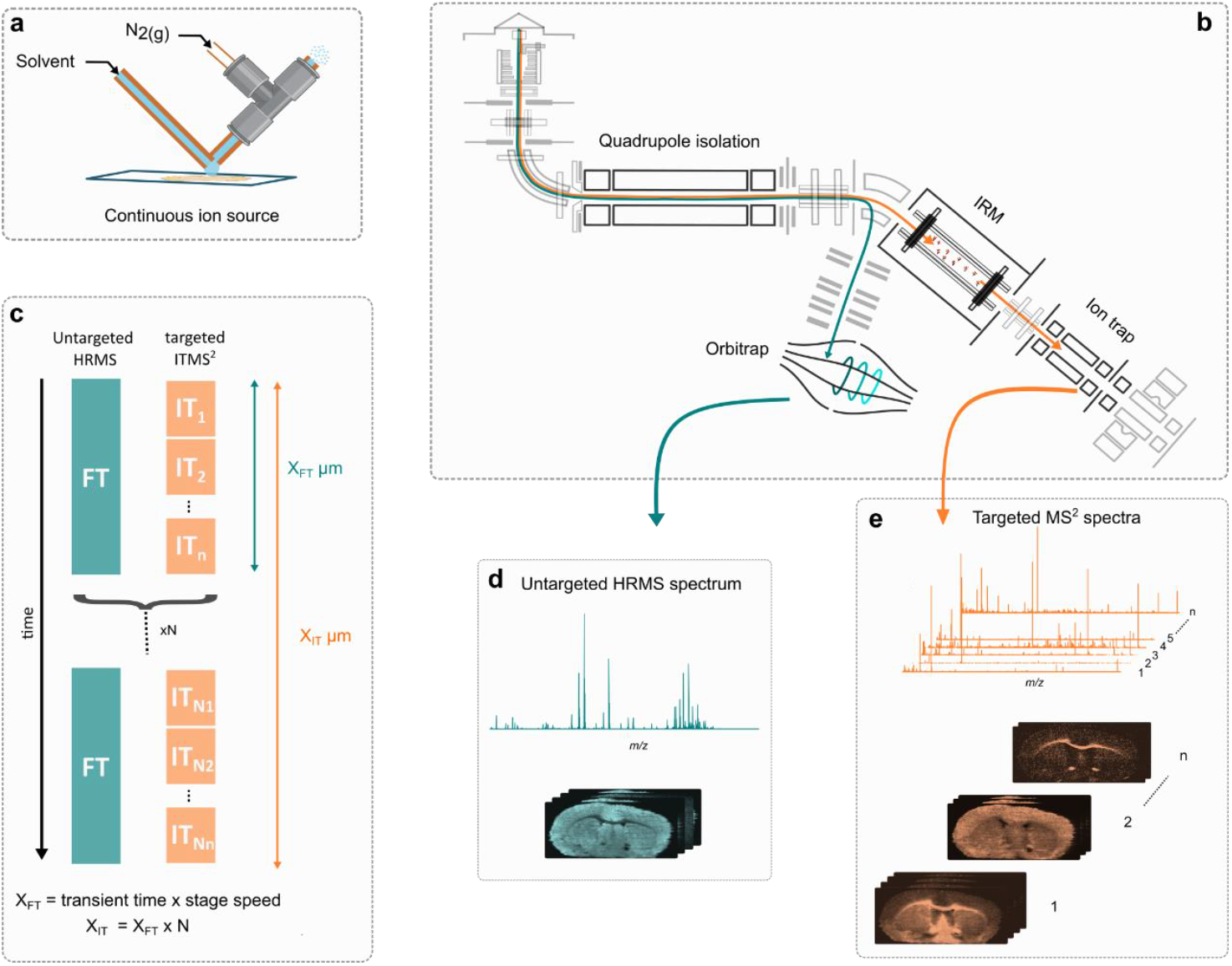
Schematic of Parallel Image Acquisition (PIA) enabling scalable MS^n^I. **a**, PA nano-DESI enables continuous sampling and ionization of molecules from tissue surfaces under ambient conditions, generating high ion flux. **b**, A tribrid mass spectrometer equipped with a quadrupole, ion trap, and Orbitrap allows simultaneous acquisition of FTMS and ITMS^2^ data, enabling PIA. **c**, Schematic of the PIA acquisition strategy: one high-resolution FTMS scan is acquired in parallel with *n* targeted ITMS^2^ scans per pixel. Cycling through *N* inclusion lists extends precursor coverage, enabling scalable MS^2^ imaging without increasing acquisition time. **d**, PIA produces untargeted, full-scan, high-mass resolution FTMS spectra with spatial information. **e**, Simultaneously acquired targeted MS^2^ spectra and corresponding product ion images allow for structural and spatial resolution of molecular species at scale.

### Decoding Complex MS^2^I Data with Spatial Similarity Networking (SSN)

Each PIA imaging experiment generates a large, multidimensional dataset in which each pixel contains one full-scan FTMS spectrum alongside 27 ITMS^2^ spectra, each corresponding to a distinct isolation window encompassing multiple unique precursors (Fig. 2a). Annotating molecules from such complex, multiplexed product ion spectra is a formidable challenge. However, a key insight is that all product ions originating from the same precursor ion exhibit identical distribution across tissue. We developed a spatial similarity networking (SSN) approach that deconvolutes MS^2^I data by leveraging the spatial correlation between product ion images. The method extracts ITMS^2^ spectra from each pixel and computes pairwise spatial similarity between product ion images across pixels, using cosine similarity or mean squared error (MSE) metrics. The results are used to populate a (dis)similarity matrix that encodes spatial relationships between product ions (2b-d). From this matrix, an undirected similarity network is then constructed, where the nodes represent individual product ions and the edges denote spatial correlation (Fig. 2e). Specifically, an unsupervised graph-based clustering that uses a connected components algorithm identifies groups of product ions sharing the same spatial pattern, without prior knowledge of precursor number, identity or fragmentation pathways. By treating each product ion image as an independent variable, SSN enables structural annotation to be directly derived from spatially coherent ion clusters. Thus, the SSN introduces spatial distributions as a third orthogonal dimension, alongside *m/z* and MS^2^ spectra, to facilitate unambiguous annotation, even in the presence of isobaric and isomeric overlap. In addition to MS^2^I, SSN enables non-targeted full-scan MSI data exploration (Supplementary Fig. 2). The SSN is integrated into our *ion-to-image* (i2i) software package, enabling broad accessibility across imaging platforms (Supplementary Fig. 3). When paired with product ion libraries (Supplementary Tables 1-3), i2i supports automatic structural annotation by matching spatially clustered networks of product ions to molecular candidates. Final identifications are validated against accurate mass, providing a robust, multidimensional framework for confident molecular annotation (Fig. 2f-g). For highly heterogeneous tissues, the SSN performance can be further enhanced by a Coherent Spatial Similarity Networking (Coherent SSN) workflow (Supplementary Fig. 4). This approach employs a patch-based image decomposition, emphasizing regional spatial coherence while suppressing localized noise, thereby yielding biologically meaningful ion clusters with enhanced robustness. Overall, the highly convoluted imaging data acquired by the multiplexed PIA approach is readily deconvoluted for annotation of targeted and non-targeted molecules in tissue.

**Fig 2.**
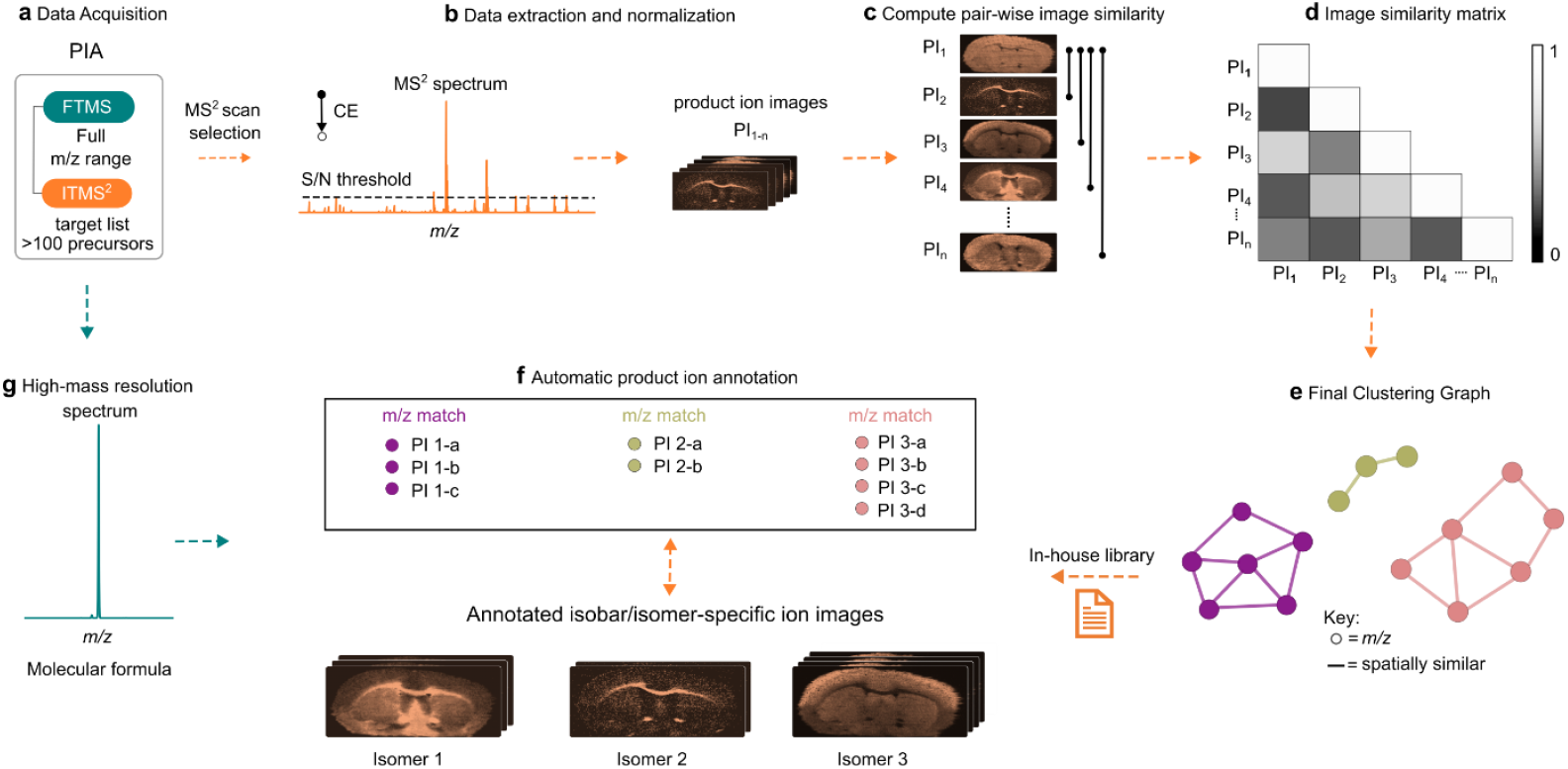
Spatial Similarity Networking (SSN) for deconvoluting complex MS^2^I data and enabling structure-resolved annotation. Product ion images from PIA (orange) are grouped by precursor isolation window and analyzed for spatial correlation using SSN. **a**, Full-scan FTMS imaging is acquired across the entire *m/z* range, while targeted MS^2^ spectra are acquired in parallel using narrow isolation windows (0.7 Da) for over 100 unique precursors. **b**, MS^2^ spectra from a selected isolation window are reconstructed into individual product ion images. **c**, Pairwise spatial similarity is computed between product ion images using either MSE or cosine similarity. **d**, Similarity matrix for precursor isolation window. **e**, Product ions are clustered using a connected components algorithm based on spatial similarity. Example output networks illustrate structurally distinct clusters arising from isomeric or isobaric precursors. **f**, Each spatially coherent cluster is matched against a curated in-house product ion library to propose molecular identities. **g**, The molecular identities are backed by high-mass resolution data. SSN leverages spatial distributions as a third dimension of evidence, complementing accurate mass and fragmentation to enhance annotation confidence.

### PIA and SSN Enable Isomer-and Isobar-Resolved Molecular Imaging and Annotation

One group of endogenous molecules with diverse isomeric structures is phospholipids, consisting of a hydrophilic phosphate head group and two hydrophobic tails derived from fatty acids. Each phospholipid class is distinguished by a specific head group, while variation in the chain length, saturation, and positional arrangement of the fatty acyl chains gives rise to a variety of isomers indistinguishable by mass spectrometry alone. Moreover, many phospholipids in complex biological mixtures exist as near-identical *m/z* (isobars) that cannot be resolved even with high mass resolution. Yet, accurate identification of these isomers and isobars is essential as they individually regulate cell and tissue functions by influencing cell membrane permeability, fluidity, and stability in addition to signalling and energy storage.^33–35^ The distribution and annotation of phospholipid isomers and isobars in mouse brain tissue were targeted in a multiplexed imaging experiment using the PIA and SSN workflow (Fig. 3a). For PIA, one FT scan with parallel scans having four looping inclusion lists (27 × 4; ±0.35 Da) were used, which resulted in 108 isolation windows and over 800,000 MS^2^ HCD spectra in a single experiment. Despite the use of narrow isolation windows (0.7 Da), co-isolated isomers and isobars still produced highly convoluted product ion spectra, resulting in multiple product ion clusters per isolation window in SSN (Supplementary Fig. 5). For example, within a single window centred at *m/z* 808.58, three discrete product ion clusters were identified with the SSN (Fig 3a). When visualizing the clustered product ion images, each group displayed distinct spatial distributions (Fig. 3b and Supplementary Fig. 6). By integrating *m/z* of the clustered product ions, molecular candidates predicted from rule-based fragmentation patterns in our in-house phospholipid library and the accurate precursors mass from FT, we annotated molecular structures down to fatty acid acyl chain composition (Fig. 3c and Supplementary Fig. 7-8). Specifically, 3 isobars [PC(36:2)+Na]^+^ (*m/z* 808.5832), [PC(38:5)+H]^+^ (*m/z* 808.5856), and [PC(O-36:3)+K]^+^ (*m/z* 808.5622) were confidently identified. While [PC(36:2)+Na]^+^ and [PC(38:5)+H]^+^ are indistinguishable in the FTMS spectrum, the distinct distributions of their product ions reveal their presence. Furthermore, [PC(36:2)+Na]^+^ was spatially deconvoluted into two structural isomers, [PC(18:1_18:1)+Na]^+^ and [PC(18:0_18:2)+Na]^+^ based on their unique product ion patterns. In another case, a single window centred at *m/z* 810.56, yielded three SSN clusters; two co-isolated isobars indistinguishable accurate mass [PC(38:4)+H]^+^ and [PC(36:1)+Na]^+^, where the latter further separated two isomeric species [PC(18:0_18:1)+Na]^+^ and [PC(16:1_20:0)+Na]^+^ each with unique spatial distributions (Fig 3d-f and Supplementary Fig 9-10). Note that a limitation of SSN can be reduced performance for low-abundant product ions due to increased signal-to-noise variability. However, the results demonstrate that the SSN successfully leverages spatial correlation among product ion images to distinguish co-isolated species that would otherwise remain unresolved, both spectrally and spatially, thereby increasing confidence in annotation and revealing molecular insights in the ion images.

**Fig 3.**
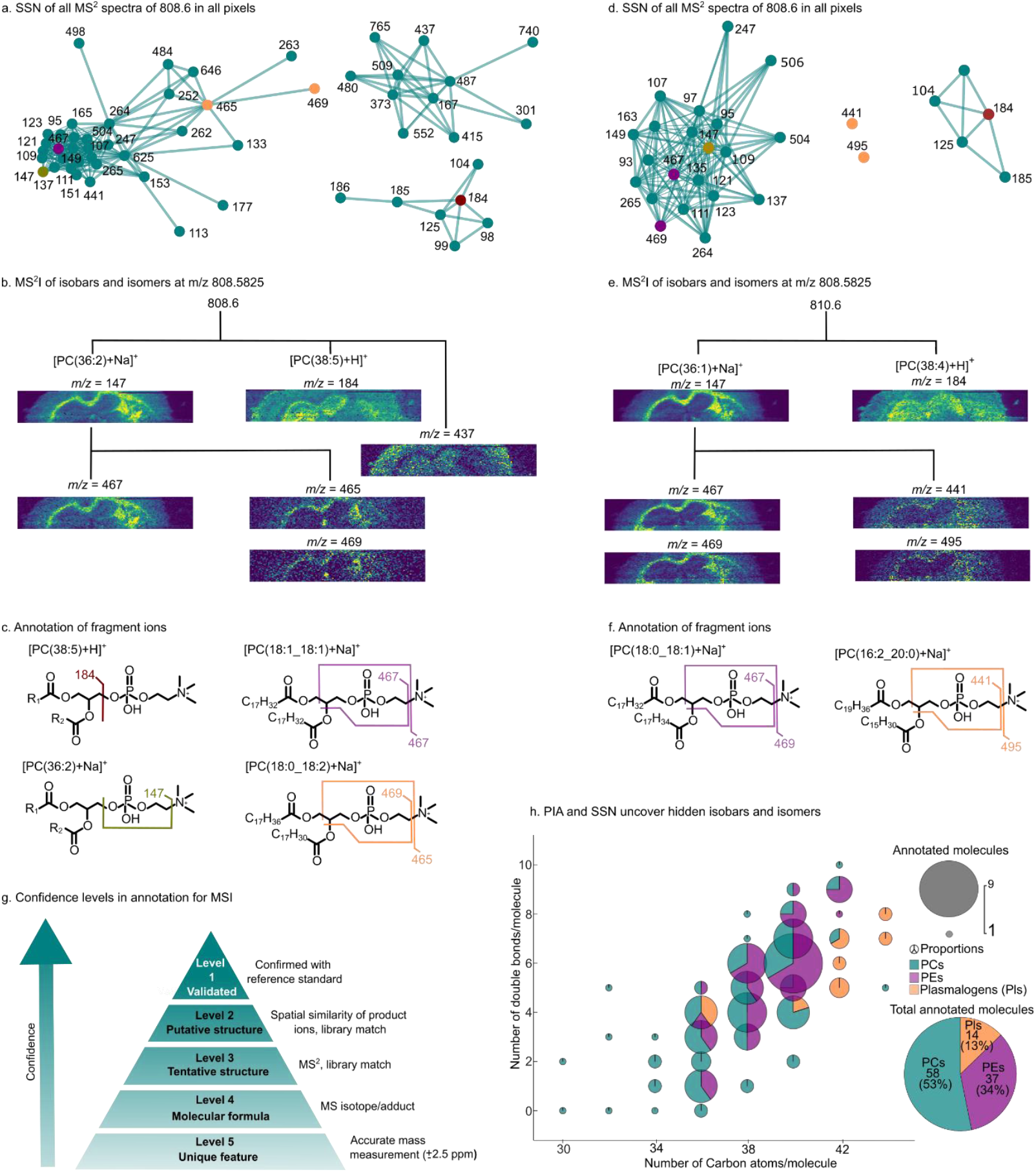
Annotation with PIA and SSN reveal molecular distributions hidden from conventional MSI. a, The SSN of product ion images derived from the fragmentation of *m/z* = 808.6 reveals three distinct clusters of product ions, each showing unique spatial distributions. This indicates the presence of multiple co-isolated precursors. **b**, The MS^2^I ion images illustrate varied molecular distributions within each cluster. **c**, Fragmentation of the head group and acyl chains of phosphatidylcholines (PCs) through HCD exposes isobars such as [PC(38:5)+H]+ and [PC(36:2)+Na]+, alongside isomers like PC(18:1/18:1) and PC(18:0/18:2).**d-f**, Fragmentation of m/z = 810.6 revealed the distribution of two isobars [PC(38:4)+H]^+^ and [PC(36:3)+Na]^+^ and two isomers PC(18:0_18:1) and PC(16:1/20:0) **g**, The suggested confidence level for annotations in MSI based on SSN analysis. **h**, bubble pie chart summarises PIA SSN-based results from 34 precursor isolation windows, identifying 134 lipid species including phosphatidylcholines (PCs), phosphatidylethanolamines (PEs), and plasmalogens (Pls). Out of these, 109 were unique phospholipid structures corresponding to 58 PCs, 37 PEs, and 14 PE plasmalogens/alkyl ethers could not be uniquely resolved by MS^1^ imaging alone, highlighting the power of SSN for resolving isobaric and isomeric complexity in situ.

Chromatographic separation prior to MS increases confidence level in annotation, as product ions originating from the same precursor co-elute, sharing the same retention time. Analogously, the SSN approach improves confidence in annotation in MSI by leveraging the co-localised spatial distributions of product ions derived from the same precursor. In this context, the spatial distribution of product ions serves as a separation dimension, alongside accurate mass and product ion spectra, to strengthen the specificity of molecular assignment. To standardise confidence levels in MSI annotation, we propose a new guideline for classifying confidence levels for annotation (Fig. 3h). This system ranges from Level 5, based solely on with accurate mass to Level 1, which necessitates confirmation against a reference standard and orthogonal validation; generally impractical at the pixel level for MSI. Within this framework, we define Level 2 confidence as requiring spatial correlation of product ions across all pixels. Such spatial co-localization links fragments to their precursor ions offering reproducibility, structural resolution, and improved specificity in annotation. This spatial-spectral concordance, which is achievable with SSN, significantly reduces the risk of false positives and misannotations, offering a robust strategy for interpreting MSI datasets and advancing molecular imaging towards more confident structural assignment.

Annotation at Level 2 was achieved in 34 targeted isolation windows using the SSN annotation workflow in combination with PIA for phospholipid imaging. Across these isolation windows, we spatially resolved and annotated 134 distinct isomeric and isobaric lipid species. These included 70 phosphatidylcholines (PCs), 44 phosphatidylethanolamines (PEs), and 20 PE-plasmalogens which were reproducibly detected across several datasets (Fig. 3g, Supplementary Tables 5–6). Notably, 75 (55%) of these species could not be resolved by FTMS imaging alone, despite the high mass resolution (*m/Δm* = 500,000 at *m/z* 200), highlighting the critical role of MS^2^I in resolving structural complexity within tissues. Ultimately, the combination of PIA and SSN uniquely identifies, distinguishes, and visualises superimposed isobaric and isomeric spatial distributions, providing a solid foundation for biological interpretations.

### Annotation and visualization of sterol lipids in human multiple sclerosis brain tissue

Sterol lipids represent a structurally diverse class of molecules, characterized by numerous isomeric structures arising from variations in unsaturation levels and hydroxylation positions.^36^ Although the role of sterols as key regulators of fundamental biological processes is well established,^37,38^ the low physiological concentrations and poor ionisation efficiency of sterols have limited their spatial analysis by MSI.^39–41^ However, we show that silver ions directly doped into the nano-DESI solvent generate argenated ions, which upon MS^2^ generate isomer-specific side-chain cleavages and product ions for sterol lipid annotation (Fig. 4a and Supplementary Fig. 11). The most abundant sterol lipid is cholesterol, which is a key constituent of cellular membranes and neuronal myelin sheets.^42^ With demyelination being a major pathological hallmark of the chronic neurodegenerative disease multiple sclerosis,^43^ mapping sterol lipids in MS brain tissue could provide insights into cholesterol clearance and disease-related pathways. Although altered levels of sterol lipids are found in cerebrospinal fluid of people with multiple sclerosis (PwMS),^44,45^ but to date their spatial distribution in brain tissue of PwMS remains elusive. PIA of a human brain tissue from a subject with multiple sclerosis, reveals the high molecular complexity of the human brain (Fig 4b and Supplementary Fig. 12). For example, within a selected isolation window of *m/z* 539.2 ± 0.35 Da, targeting dihydroxycholestenoic acid, the FTMS spectrum shows co-isolated precursor ions each with distinct spatial distributions over the tissue contributing to the MS^2^ spectrum. Despite this complexity, SSN effectively deconvoluted the MS^2^ spectra, clustering the product ions into six distinct groups (Fig. 4c). One cluster, comprising three product ions with highly similar spatial distributions, was annotated as dihydroxycholestenoic acid (ST5) based on known diagnostic product ions (Fig. 4d and Supplementary Table 7). The combination of PIA and SSN thereby provides annotation beyond accurate mass and visualization with isomeric precision. In total, the distribution of cholesterol and six sterol lipids was successfully mapped in human brain tissue using the PIA and SSN workflow (Fig. 4e and Supplementary Figs. 13-15). Beyond demyelination, a second key pathological feature of multiple sclerosis is blood–brain barrier (BBB) disruption.^46^ By combining traditional immunohistochemical staining techniques with higher resolution PIA, images generated over a blood vessel revealed leakage of the bile acid 24-homodeoxycholic acid (ST7) (Fig. 4f). Changes in bile acid metabolism have recently been shown to change in the course of multiple sclerosis development.^35^ Since these sterols are not biosynthesized within the brain, our results strongly support the occurrence of BBB disruption in multiple sclerosis pathology.^47^ Overall, by combining enhanced ionization strategies with PIA and SSN, the spatially resolved mapping of structurally complex, low-abundance, understudied sterol lipids can provide insights into disease-related molecular processes in situ.

**Fig 4.**
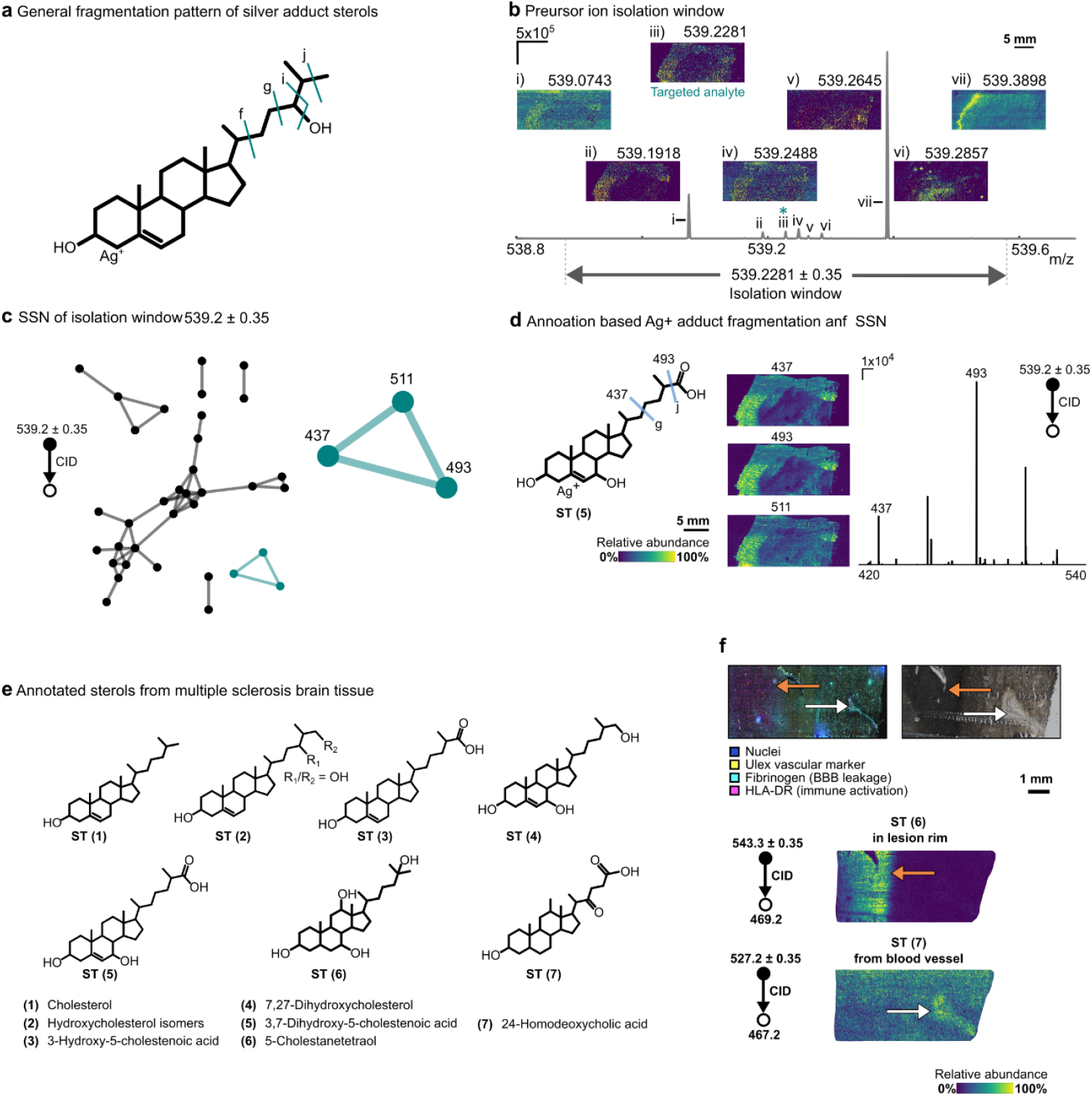
Detection and annotation of sterol lipids in human multiple sclerosis brain tissue. **a**, Sterols, ionized as silver adducts, exhibit consistent fragmentation in their side chains as illustrated for 24-hydroxycholesterol. **b**, An FTMS spectrum detailing the narrow isolation window (0.7 Da) of ions selected for ITMS^2^ with corresponding high-mass resolution precursor ion images. **c**, SSN clusters all product ion images based on spatial similarity. The network highlighted in green corresponded to the side chain fragmentation of a sterol lipid fragmented as silver adduct. **d**, Side chain fragmentation matches with 3, 7-dichydroxycholestenoic acid, and the three clustered product ions show very similar distributions, indicating they arise from the same molecule. **e**, The structure of cholesterol and six annotated oxysterol species from human multiple sclerosis brain tissue. **f**, A multimodal study combining PIA, SSN and histological staining reveals infiltration of homodeoxycholic acid to a white matter lesion through a BBB leakage site. Representative image of UEA-I, Fibrinogen and HLA-DR immunoreactivity and optical image with corresponding images of diagnostic product ions for cholestanetetraol (ST 6) and 24-homodeoxycholic acid (ST 7), white arrow: BBB leakage site. CID, collision-induced dissociation.

### Multimodal oxysterol landscape is altered in multiple sclerosis

The oxidation of cholesterol in multiple sclerosis may be directly linked to lesion formation and disease progression.^44^ White matter brain tissue from three non-neurological control (NNC) and five PwMS were imaged in a multimodal setting using both silver doped PA nano-DESI MSI, PIA, and immunohistochemical techniques (Supplementary Fig 16). Following, the stage of demyelination was determined using proteolipid protein (PLP) expression and categorized as normal PLP, diffuse PLP, and absent PLP, where the absent PLP indicate most severe demyelination. The spatial information was transferred to the ion images where regions of interest (ROIs) were defined, stratified into normal PLP (n=19), diffuse PLP (n=20), and absent PLP (n=16) as described in the Methods section, and data on sterol lipids ST1, ST3, ST4 and ST7 were extracted (Fig 5a). The multimodal results show a significant 2-fold depletion of cholesterol that correlate with demyelinated regions. Additionally, the ST4 and ST7 were significantly increased (∼2-fold) in regions exhibiting diffuse and absent PLP compared to areas with normal PLP, while an increasing trend was observed for ST2 and ST3, which is indicated as being neurotoxic.^48^ Looking at the overall oxysterol landscape, we observed a considerable differentiation in the grouping based on the subjects being NNC or PwMS (Fig. 5b-d, Supplementary Fig. S17), with the dominant sterol in NNC and PwMS samples being cholesterol and hydroxycholesterol, respectively. Furthermore, normal PLP ROIs were only slightly separated from diffuse and absent PLP ROIs. This indicates that the decreased cholesterol level and elevated cholesterol oxidation may be a combination of myelin degradation and other effects, including local inflammatory and immune response. Overall, we find an interchange between cholesterol and oxidized cholesterol species in the demyelination process in multiple sclerosis and an overall elevated cholesterol oxidation in PwMS brain tissues, which highlights the importance of spatially resolved isomer specific data and lay the basis for further investigations.

**Fig 5.**
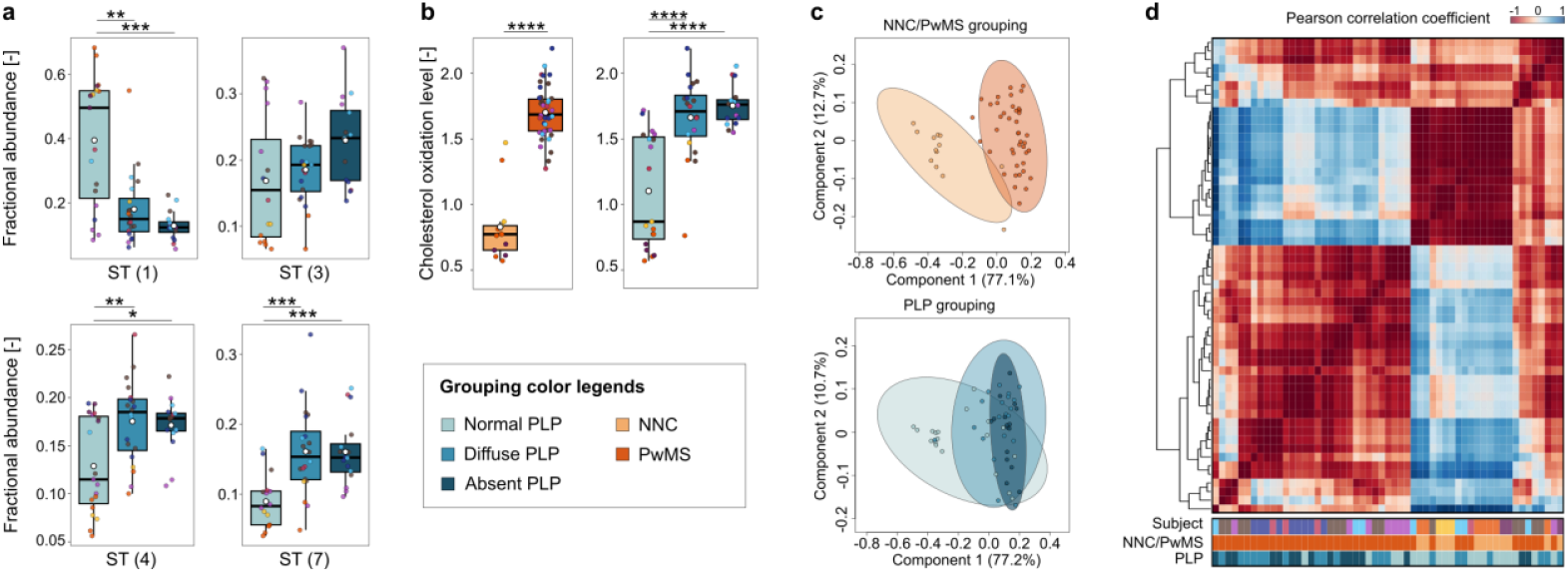
The landscape of cholesterol and cholesterol oxidation products is altered in multiple sclerosis. **a**, Boxplots represent the fractional abundance of significantly decreased cholesterol (ST 1) and increased oxysterol (ST 3, ST 4, ST 7) species. **b**, Levels of sterol oxidation significantly increases in diffuse and absent PLP category compared to normal PLP and is higher in PwMS compared to NNCs. For calculations see Methods section. Point colors indicate individual donors as per Supplementary Table 8, black line denotes the median and the white point the mean. **c**, PLS-DA analysis of sample grouping shows considerable differences according donor type (NNC vs PwMS) and slight separation of absent PLP from normal and diffuse PLP groups based on oxysterol profiling of ROIs. **d**, Pearson correlation heatmap of ROIs reinforces the PLS observations by showing primary clustering based on donor type and secondary clustering based on PLP grouping. PLP, proteolipidprotein staining of myelin; NNC, non-neurological control; PwMS, people with multiple sclerosis. (*:p<0.05, **:p<0.01, ***:p<0.001, ****:p<0.0001).

## Conclusion

We present a novel approach for structurally resolved MSI by combining PIA and SSN in a powerful platform for high-precision spatial omics. PIA and SSN facilitates both extensive molecular coverage from MS^2^I and detailed molecular structure analysis based on spatial distribution, with the SSN seamlessly integrated into our i2i software. In particular, we show the distribution and detailed molecular structure characterization of isomeric and isobaric phospholipids mouse brain tissue in addition to the much less abundant oxysterol in human brain tissue. Additionally, we suggest new annotation levels for strengthening reported annotation by MSI, increasing validity for the entire MSI community. In the context of multiple sclerosis, we observed strong indications of increased sterol leakage into the brain parenchyma and the implication of cholesterol oxidation in demyelinated multiple sclerosis. While our applications primarily highlight metabolomics, the PIA and SSN strategy is versatile and applicable to proteomics, glycomics and pharmacokinetics. Moreover, coupling PIA and SSN with higher spatial or tandem mass resolution instrumentation,^49^ or reactive chemistry workflows,^50,51^ could further deepen structural characterization. Ultimately, PIA and SSN pioneer the integration of large-scale information-rich molecular distributions with enhanced structural characterization, setting a new benchmark in spatial omics.

## Methods

### Sample selection for model experiments

Mouse brain (8–10 weeks old male C57BL/6) was purchased from Creative Biolabs (Shirley, NY, USA) and cut to a thickness of 10 µm using a cryo-microtome (Leica Microsystems, Wetzlar, Germany). The sections were thaw-mounted onto regular microscope glass slides and stored at -80 °C prior to analysis.

### Sample selection for analysis of human tissue samples

Freshly frozen post-mortem human brain samples were obtained from the Netherlands Brain Bank from 5 clinically diagnosed MS patients (mean age at death: 65 years) and 3 non-neurological controls (mean age at death: 66 years). Tissue blocks were cut into 10 µm sections and stored at -80 °C until further use. All donors or their next of kin provided fully informed consent for autopsy and use of material for research from the Netherlands Brain Bank under ethical approval by the Medical Ethics Committee of the Free University Medical Center in Amsterdam (2009/148), project number 1,127. Donor details are shown in Supplementary Table 8.

### Parallel image acquisition using PA nano-DESI

Samples were gently thawed at room temperature under constant airflow before analysis. Mouse brain sections were analyzed without any sample preparation, human multiple sclerosis tissue sections were dipped six times in deionized water (Milli-Q, 18.2 MΩ) for one minute each to remove excess salts. For MSI, the samples were placed on an XYZ linear motor stage (Zaber Technologies Inc., Vancouver, BC), controlled via a custom-designed LABVIEW program.^52^

The PA nano-DESI was set up as previously described,^28^ using 150/50 µm o.d./i.d. fused silica capillaries (Genetek, Sweden). Briefly, the primary capillary, connected to a syringe containing the nano-DESI solvent, was coupled to a pneumatically assisted secondary capillary positioned at ∼90 degrees. The extraction solvent was delivered through the primary capillary via a syringe pump (Legato 180, KD Scientific, Holliston, USA), with the ESI voltage applied directly to the syringe needle. The probe and sample positioning were monitored using long-working-distance digital optical microscopes (Dino-Lite, USA). For all experiments the internal Detailed PA nano-DESI parameters are provided in Supplementary Table 9.

Mass spectrometry was performed in positive ion mode on an Orbitrap IQ-X (Thermo Fisher Scientific, San Jose, CA, USA) using high-resolution Orbitrap (FTMS) MS^1^ and ion trap (ITMS) MS^2^ modes, or FTMS^2^ for accurate mass confirmation. For PIA, ions were first selected for a full FTMS scan and subsequently for several parallel MS^2^ scans in the ion trap during the FTMS transition time.^53^ The PIA methodology was tested across five different experimental conditions (varying extraction solvents and targeted analytes) to demonstrate its versatility (Supplementary Table 6). Detailed mass spectrometry acquisition parameters for each experiment are provided in Supplementary Table 9.

Figures showing data and metadata from the experiments were exported from FreeStyle 1.8 SP2, v1.8.63.0. Ion images were generated using our in-house software tool, i2i,^54^ from .mzML data converted via MSConvertGUI (ProteoWizard, v3.0.22285) from .RAW files. The use of. mzML format facilitates a vendor-neutral use of the software. Ion images of the MS^2^ data was created by imputing the scans on a master grid generated from the FTMS scans where the specific MS^2^ spectra were given the same coordinates as the closest FTMS master scan.

### Spatial similarity network analysis

The SSN is computed based on the intensities of all identified *m/z* values in all pixels. First, the user selects the scan filter, scan type, and search parameters. Although the GUI provides suggested pre-set parameters for ppm tolerance and min and max intensities, these may need to be altered based on the experiment. Secondly, the user selects if the SSN should be computed using either the sum squared error (SSE) or the cosine distance metric.

When the user clicks the run button, a non-targeted search, based on the user-defined parameters, is conducted to identify *m/z* values that will be considered for the SSN. The algorithm for the search is identical to the algorithm in the non-targeted tab of i2i.^54^ Following, one of the *m/z* values is selected as an *anchor ion image* and the distance metric is calculated for all identified *m/z* values against the anchor ion image. To *normalize* the response of different ions, all compared *m/z* values are first scaled to the 99^th^ percentile of intensity within their respective ion image. This allows for comparison across *m/z* values despite differences in absolute intensities. Subsequently, the anchor ion image is exchanged to ensure that all ion images of identified *m/z* values are compared with each other.

The equation used to calculate the SSN based on SSE and cosine distance is shown in Eq. 1 and Eq. 2, respectively. The x and y are intensities of the respective *m/z* in a given pixel, the *x*_*anchor*_ is the matrix of the anchor ion image and *y*_*i*_ is the matrix of the ion image for the *i*^th^ ion.

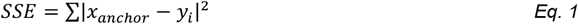

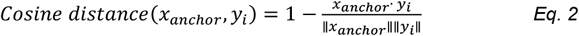

The computational process creates a lower triangular correlation matrix with values corresponding to a similarity index (SI). The use of a lower triangular matrix saves memory usage and computation time during the search process. The ion images with high SI, as set by the user as the error limit, are grouped based on the user-selected percentile of the similarity index values to be included. For example, by selecting 0.1, only 10% of the lowest SI values (i.e. the SI values with the highest correlation) are used to generate a filtered correlation matrix. The error limit has to be optimized for each experiment with respect to the complexity of the given scan event. A graph in MATLAB is then created with the filtered correlation matrix, with each group in the correlation matrix corresponding to a connected component of the graph. Following, the function conncomp is used to extract the spatially correlated *m/z* values of the groups. The results are shown as clusters of *m/z* values with similar distributions, termed the SSN.

To interrogate the SSN, the user can select groups and their *m/z* values in the list box next to the displayed SSN. Alternatively, by opening the SSN as a plotted figure, the user can hover over the displayed SSN to view information on each node in real-time and save the plot. The number of individual SSN groups is ordered by size, with Group 1 being the largest. The graph layout, including the length of the lines within each SSN group, is automatically set by MATLAB and has no value for the interpretation of the SSN. For automatic annotation of the *m/z* values in the SSN based on user-selected ppm difference, the user can upload a database containing *m/z* values and known names of the corresponding ions. The annotated ions are identified in the list box by a green dot next to the *m/z* value. All SSN calculations and interrogation are supported by a GUI integrated into our i2i application (github: LanekoffLab),^54^ and the functionalities are detailed in Supplementary Fig. 3.

### Immunostainings and tissue characterization

Fluorescent immunostainings were performed as previously published.^47^ Briefly, frozen human tissue slides were dried at room temperature, fixed with 4% PFA at room temperature for 10 min, washed with PBS and incubated with blocking solution (10% normal species serum (NSS) in 0.05% Tween20 (Sigma-Aldrich)) for 30 min. Primary antibodies for PLP (1:300; Serotec, MCA839G), UEA-I (1:1000, Vector Labs B-1065), HLA-DR (1:500; Hybridoma) and Fibrinogen-FITC (1:300, Dako, F0111) were diluted in 1% NSS in 0.05% Tween20-PBS and applied on the slides overnight at 4 °C. Tissue sections were washed in PBS and incubated for 1 h with Alexa fluorophore-conjugated secondary antibody. Nuclear staining was performed using Hoechst fluorescent DNA stain (33258, Thermo Fisher Scientific) for 1 min. Images were acquired as a 20x overview scan at the Olympus VS200 slide scanner. Following, visual characterization was performed using QuPath software (version 0.4.4) to aid classification based on full (normal), diffuse (dirty appearing white matter) and absent (lesion) PLP immunoreactivity.^55^

### Regions of interest selection and data extraction

Tissue samples from three non-neurological control (NNC) and five people with multiple sclerosis (PwMS) were analyzed using one brain tissue section for each subject. A multimodal approach was used to select ROIs unambiguously. In particular, a region had to fulfil two criteria to be considered as an ROI: i) have a morphologically well-defined border from surrounding areas based on PLP and HLA-DR staining, ii) have a well-defined chemical profile based on ion images acquired using PA nano-DESI MSI. Morphologically distinct areas were defined according to the section “Immunostainings and tissue characterization” and stratified into normal, diffuse and absent PLP categories. (Extended Data Fig. 7). Chemically distinct areas were defined by segmenting the PA nano-DESI MSI ion images based on the spatial distribution of monoacylglycerol, phosphatidylcholine, phosphatidylethanolamine, prostaglandin, free fatty acid and sterol lipid classes. In total, 55 ROIs were identified: 19 normal PLP, 20 diffuse PLP, and 16 absent PLP (Supplementary Table 8).

To compare the abundance of cholesterol and oxidized cholesterol products, average pixel intensities were extracted from each ROI using the i2i software^54^ and the fractional abundance (FA) was calculated (Eq 3). In Eq. 3, the FA is calculated for each oxysterol based on its intensity over the sum of intensities of cholesterol (ST1) and all selected oxysterols (ST3, ST4, ST7).

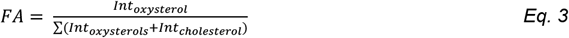

Subsequently, the cholesterol oxidation level was defined according to Eq 4, where FA is the fractional abundance of the molecule, and ON is defined by the additional oxygens in the sum-composition formula compared to cholesterol (0 for ST1, 1 for ST2, 2 for ST3 and ST4, and 4 for ST7).

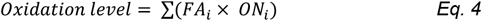

## Statistical analysis

Statistical analysis was performed, and boxplots were created using R v4.3.2 in RStudio.^56^ The normality of the data was tested using the Shapiro-Wilk test. The pilot study cohort was balanced for age and post-mortem delay, and the covariate significance of the distinct patient was tested using a Kruskal-Wallis test. Since it was significant in some cases and not significant in others, we decided to perform non-parametric one-way testing but denote each donor by color codes as per Supplementary Table 8. In multi-group comparisons, the Kruskal-Wallis test was performed, followed by Dunn’s post-test, where a significant difference was indicated. For the two-group comparisons, the Wilcoxon test was used. FDR correction was performed in all analyses using the Benjamini-Hochberg method. Significant differences are indicated in the plots as follows. *:p<0.05, **:p<0.01, ***:p<0.001, ****:p<0.0001. The results of statistical tests are detailed in Supplementary Table 10. In the boxplots, the black line denotes the median, the white dot the mean, the top and bottom edges of the box the interquartile range, and the whiskers extend to the minimum and maximum data points.

Hierarchical clustering-based heatmaps and Pearson correlation heatmaps and PLS-DA plots were created using MetaboAnalyst 6.0.^57^ Missing data points were imputed by LOD (1/5 of the lowest intensity). For hierarchical clustering, values were Z-scored, Euclidean distance measure was set, and autoscaling was set based on samples.

## Data availability

Data has been uploaded to figshare

## Code availability

The source code and the compiled version of the software will be made available upon acceptance of the manuscript at https://github.com/LanekoffLab

## Acknowledgements

We thank the donors and their families for making the tissue available in the biobank and Dr. Johan Lillja for his contributions to the initial work and fruitful discussions.

## Funding

Funding for this work was provided by the Swedish Research Council (2022-06628 and 2023-03384 to I.L.) and the European Union (ERC, 101041224 – X CELL to I.L.). Views and opinions expressed are, however, those of the author(s) only and do not necessarily reflect those of the European Union or the European Research Council. Neither the European Union nor the granting authority can be held responsible for them. This work was also supported by a grant from the Dutch Research Council (NWO Vidi grant 91719305 to G.K.).

## Author contributions

I.L. conceived the study. R.M. developed the software. V.V.S and G.T. performed mass spectrometry imaging experiments, analyzed and visualized the data. G.K. provided samples, C.H. performed immunostaining experiments and G.T. and C.H. performed tissue classification. I.L., V.V.S., and G.T. wrote the original manuscript. All authors contributed to reviewing and correcting the manuscript and gave consent to the final version.

## Competing interest

The authors declare no competing interests.

## Notes

### Competing Interest Statement

The authors have declared no competing interest.

### Summary of Updates

The paper has been reworked into a full paper format from the previous communication format.

