## Supplementary information for "Next-level high-precision spatial omics enabled through parallel image acquisition and spatial similarity networking"

**Prof. Ingela Lanekoff**

Dept. of Chemistry-BMC (576)

Uppsala University

751 23 Uppsala

Sweden

Supplementary Figure 8. Fragmentation patterns of phosphotidylcholine (PCs) and phosphotidylethanolamine (PEs) as sodiated adducts in HCD. . ....................................................................................11

Supplementary Figure 9. Product ion image networks from PIA SSN. . ..........................................................12

Supplementary Figure 10. SSN deconvolutes product ion mass spectra and facilitates annotation of isobars and isomers. .........................................................................................................................13

Supplementary Figure 11. Fragmentation of argenated adducts of hydroxycholesterols. . ..............................14

Supplementary Figure 12. The isolation window contains multiple precursors ions. ......................................15

Supplementary Figure 13. The SSN deconvolutes complex $ITMS^2$ spectra produced by the fragmentation of co-isolated precursor ions. . .................................................................................................16

Supplementary Figure 14. Detected oxysterol species from human multiple sclerosis tissue by sequential $FTMS^2I$. . .........................................................................................................................17

Supplementary Figure 15. Metabolic pathway for biosynthesis of bio acids from cholesterol. .......................18

Supplementary Figure 16. Multimodal imaging of cholesterol and oxidized cholesterol products in human white matter brain tissue sections. ..............................................................................................19

Supplementary Figure 17. Differentiation and clustering of brain tissue regions based on their oxysterol profiles. . .........................................................................................................................20

### List of supplementary tables

**Supplementary Table 1** fragmentation library for PE lipids based on characteristic and diagnostic product ions of $[M+Na]^+$ parent ions.

**Supplementary Table 2** Fragmentation library for PE plasmalogens based on characteristic and diagnostic product ions of $[M+Na]^+$ parent ions.

**Supplementary Table 3** Fragmentation library for PC lipids based on characteristic and diagnostic product ions of $[M+Na]^+$ parent ions.

**Supplementary Table 4** Tentative annotations of targeted mass channels based on accurate mass match.

**Supplementary Table 5** Acyl chain-specific annotation (after HCD fragmentation of sodiated adducts) of glycerophospholipid species in mouse brain tissue using PIA SSN

**Supplementary Table 6** Validation of annotated glycerophospholipid species in mouse brain tissue using PIA SSN with different experimental settings. (n.t.: non-targeted and no overlap with targeted isolation windows, n.d.: characteristic product ion not detected)

**Supplementary Table 7** List of diagnostic product ions (DPIs) for cholesterol and the investigated oxidized cholesterol metabolites.

**Supplementary Table 8** Medical details of human subject samples used for the multimodal characterization of cholesterol oxidation in the human multiple sclerosis brain.

**Supplementary Table 9** Detailed parameter list for PA nano-DESI MSI including full target lists for all the experiments performed using PIA SSN.

**Supplementary Table 10** Results of statistical tests during the characterization of cholesterol oxidation in the human multiple sclerosis brain.

The supplementary tables are supplied as one separate Excel file.

### Supplementary figures

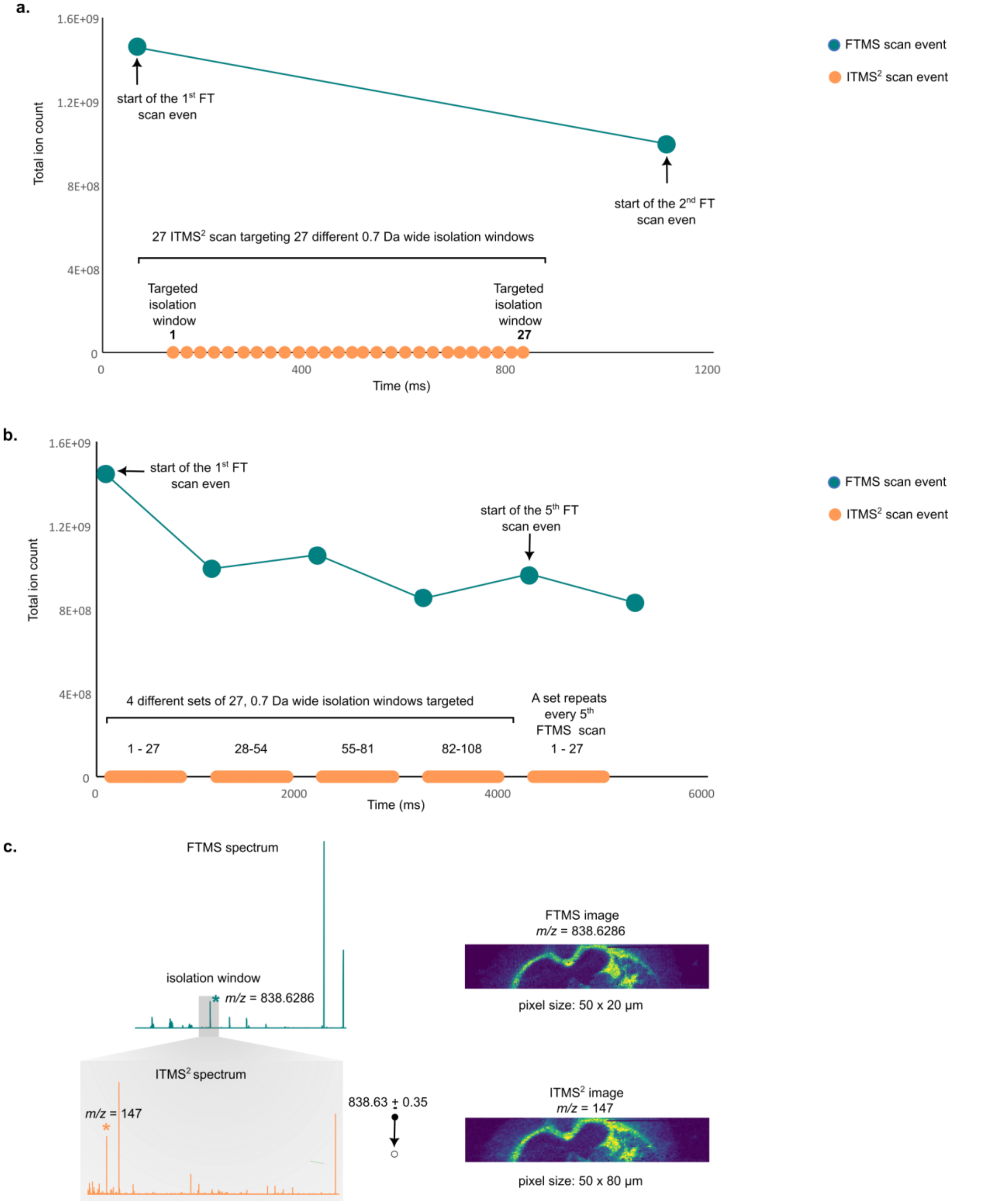

**Supplementary Figure 1. Parallel image acquisition (PIA) enables multiplexed MS² imaging without compromising acquisition time. a,** Total ion current (TIC) plotted against time illustrates the sequence of events during one complete FTMS scan at resolution 500K (m/z =200) acquired using PIA. PIA enables sequential fragmentation of 27 targeted isolation windows (0.7 Da width) using ITMS between two high-resolution FTMS scans. This allows $MS^2$ spectra to be acquired in parallel with full scan MS. **b,** PIA is scalable by looping through multiple inclusion lists. Here, four sets of 27 isolation windows (totaling 108) are fragmented between 4 FTMS scans, significantly increasing molecular coverage. **c**, Looping increases the pixel size of $ITMS^2$ data by a factor equal to the number of loops (e.g., 4× for four lists) while FTMS pixels remain unchanged. For example, the ion image of a mouse brain section with FTMS for precursor ion at *m/z* = 838.6286 has **a** pixel size of 50 x 20 µm, while its corresponding product ion in $ITMS^2$ at *m/z* = 147 has a pixel size of 50 x 80 µm.

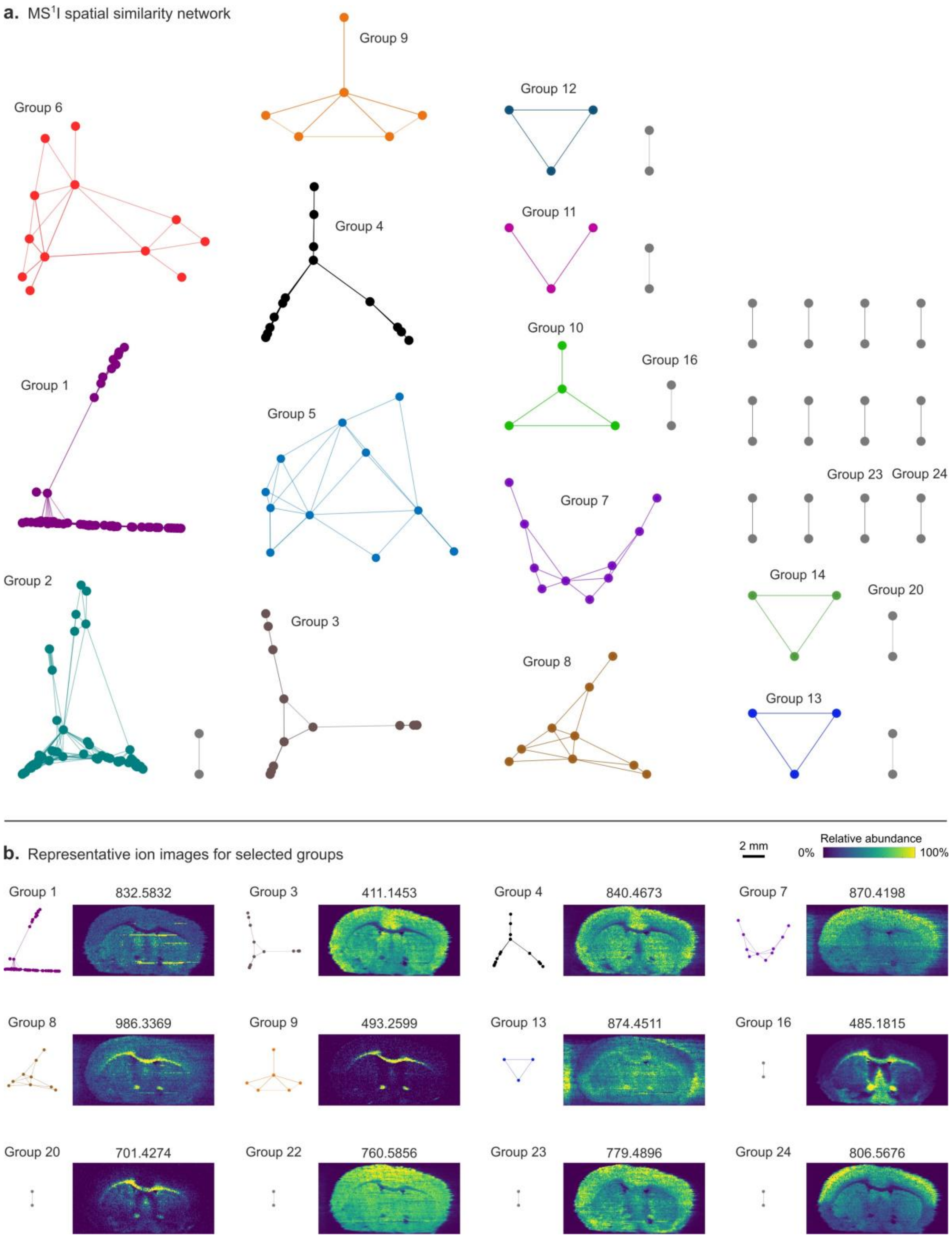

**Supplementary Figure 2. The SSN tool enables non-targeted data exploration of $MS^1$I data.** The SSN is not limited to using $MS^2$ data and product ions. On the contrary, SSN also rapidly clusters molecular ions with similar distribution, despite the increased number of available *m/z* values in the 200-2000 range. The SSN for a $MS^1$I data set displays a large complexity of spatial distributions (a). Altogether, the network analysis of 272 *m/z* values was performed, where all had at least one edge connecting to the node of another *m/z* feature. The parameters were 5 ppm mass tolerance, 0.15 error limit, 15000 intensity limit, MSE distance metric. Further 760 *m/z* values represented by disconnected nodes were discarded from the analysis. Representative ion images of separated groups using the SSN network (b) show the variety of spatial distributions of mouse brain lipids. All ion images were normalized to TIC, and intensities were scaled to the 99$^{th}$ percentile. For experimental details see Dataset 4 in Supplementary Table 9.

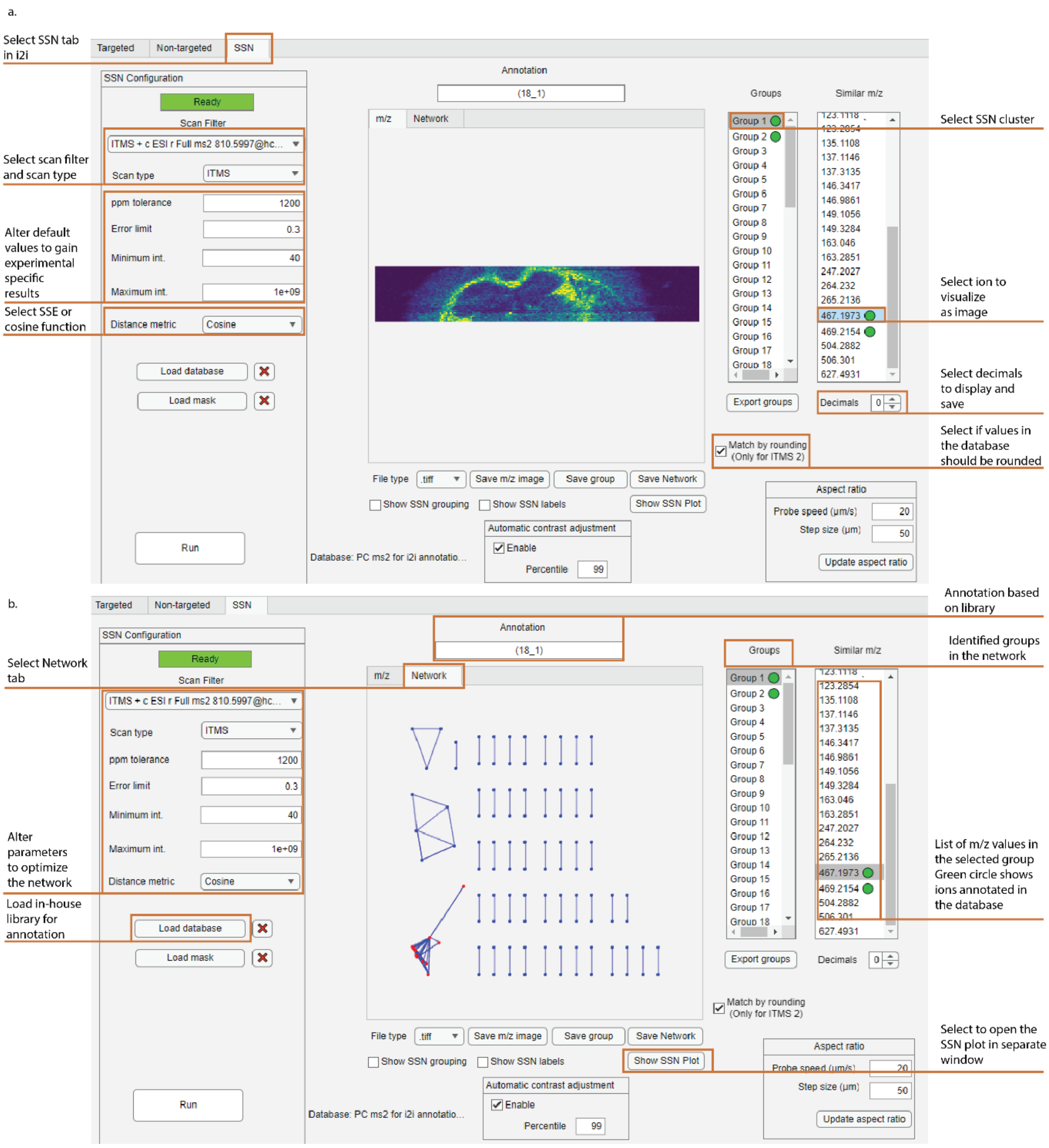

**Supplementary Figure 3. GUI of the new SSN tab in i2i.** The operator selects the scan filter and scan type based on the loaded data, along with the specified parameters. Default parameters are suggested, although optimization may be required for each scan filter depending on the complexity and intensity of the detected ions (a). The SSN automatically clusters ions in the selected scan (here, FTMS, $FTMS^2$ or $ITMS^2$) into networks based on either the sum of squared error (SSE) or the cosine spatial similarity metric of the ion distributions. The network group and *m/z* of the ion image to be displayed are selected together with the relevant number of decimals or rounding, depending on the mass resolution. The product ion image in the *m/z* tab shows the acyl chain 18:0 (*m/z* 467.2 of $[PC(36:1)+Na]^+$ (*m/z* 810.2) with the annotated ions marked with a green dot. In the network tab, the resulting SSN is displayed (b). The network shows nodes of *m/z* values that are clustered based on their spatial distributions. The number of individual SSN groups is ordered by size, with Group 1 being the largest. The graph layout, including the length of the lines within each SSN group, is automatically set by MATLAB and has no value for the interpretation of the SSN. To interrogate the SSN, the user can select groups and their *m/z* values in the list box next to the displayed SSN. Alternatively, by opening the SSN as a new window, the user can hover over the displayed SSN to view information on each node in real-time and save the plot. For the automatic annotation of m/z values in the SSN, based on ppm differences or rounding, the user can upload a database containing m/z values and the corresponding known names of the ions for matching (Supplementary Tables 1-4, 7). For experimental details, see Dataset 1 in Supplementary Table 9.

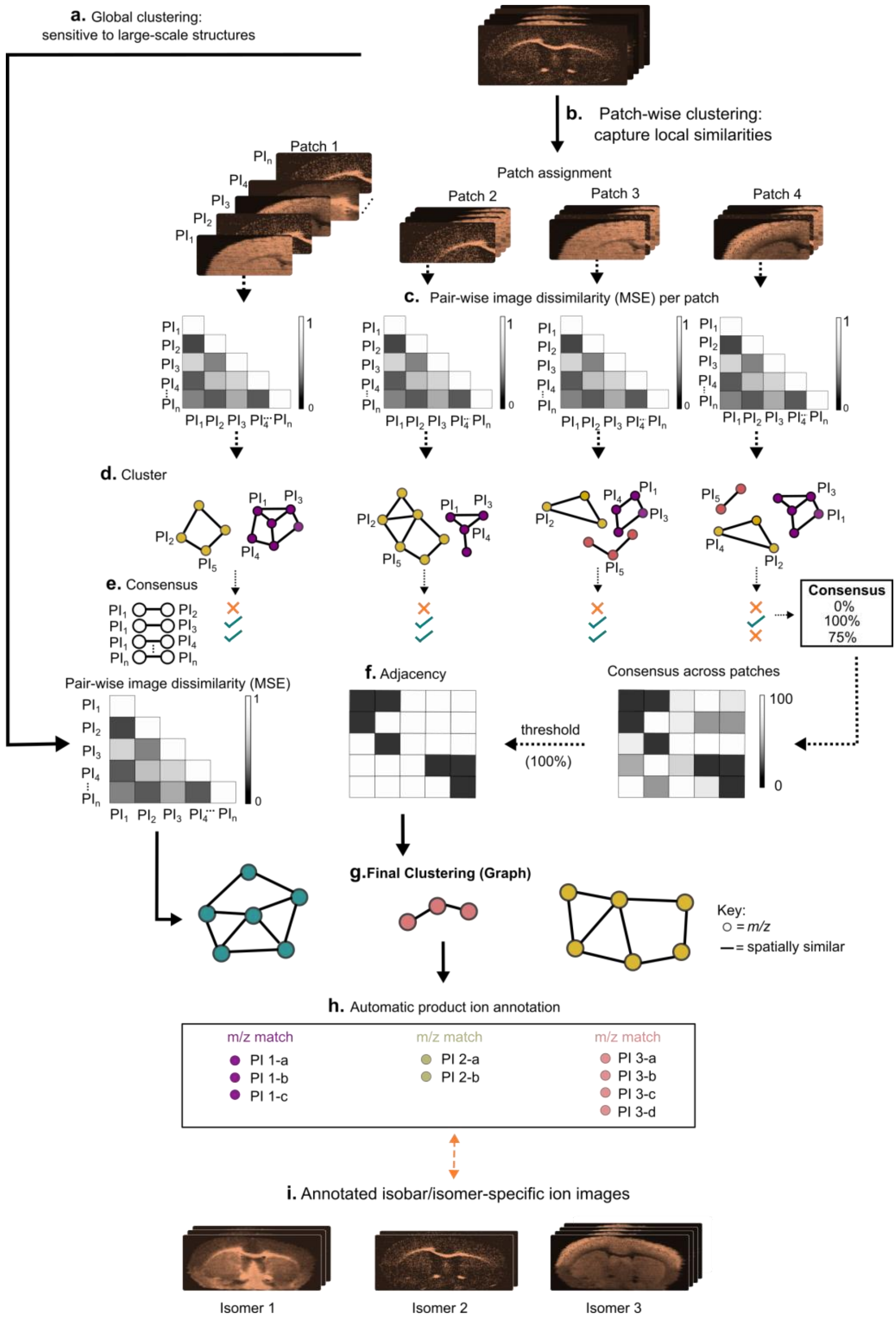

**Supplementary Figure 4. Schematics of patch-wise coherent spatial similarity networking (coherent SSN)** **a,** Global SSN network as described in Figure 2. **b-I**, patch-wise clustering workflow. b, image is divided in two equal-sized patched where the size of the patches could be 4, 9 or 16. c, Pairwise spatial similarity is computed between product ion images using either MSE or cosine similarity. d, Similarity matrix for precursor isolation window. d, Product ions in each patch are clustered using a connected components algorithm based on spatial similarity. e, A pair-wise consensus is taken for all ion images across clusters arising from different patches, resulting in a consensus matrix. For example, if a pair of ion images is connected across all clusters, then the consensus for that pair is 100%. f, an adjacency matrix is created from the consensus matrix by filtering out ion image pairs above a specified threshold. If the threshold is 100% the ion image pairs that are connected in all clusters are filtered out. **g,** Product ions are finally clustered using the connected components algorithm based on the adjacency matrix, and thus, coherent spatial similarity across patches. **h-i,** Each spatially coherent cluster is matched against a curated in-house product ion library to propose molecular identities.

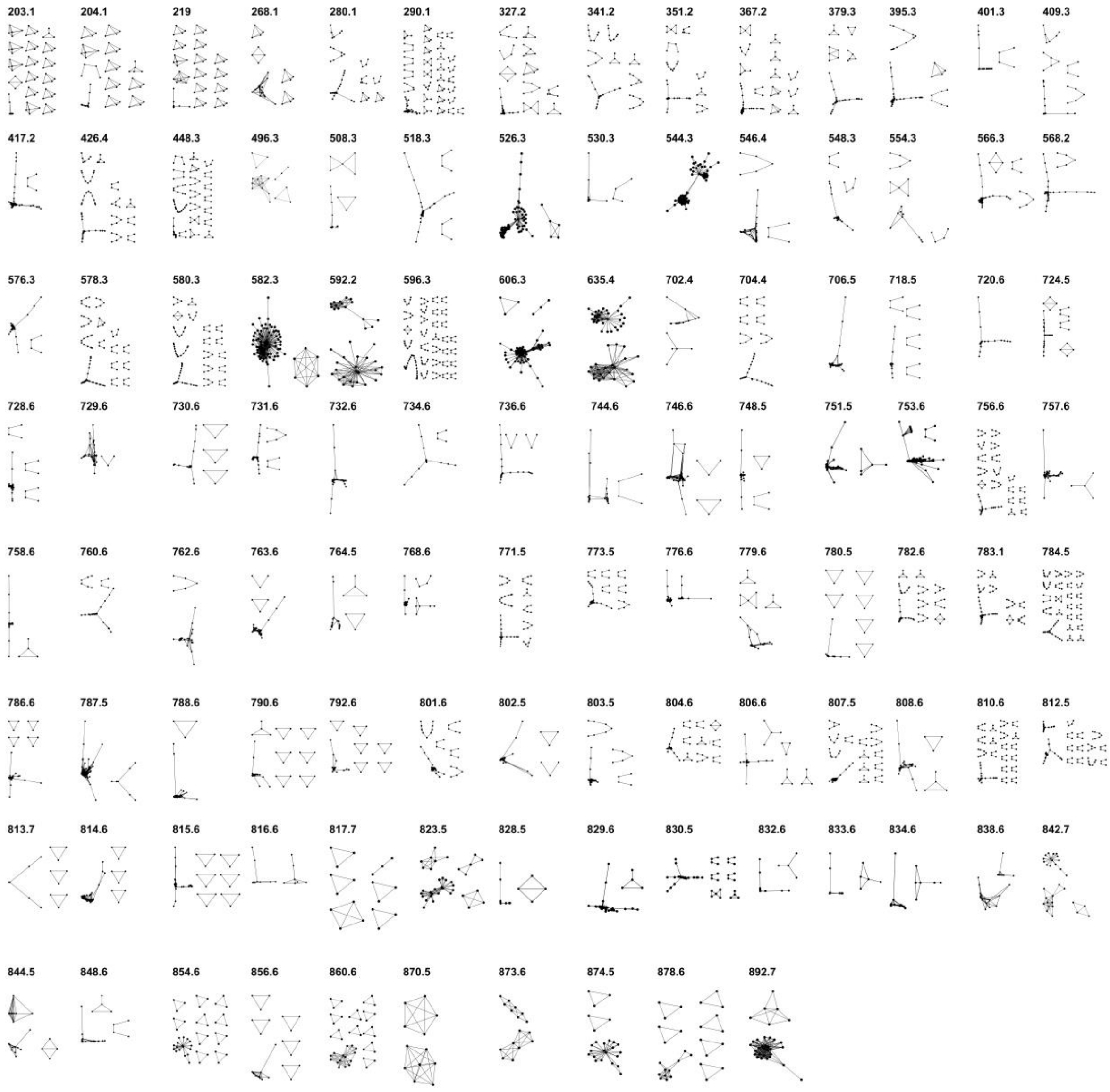

**Supplementary Figure 5. SSN of 108 targeted mass channels show a great versatility of spatial distributions.** Data from PIA (N=4, n=27) were used for the SSN with the parameters being: 800 ppm tolerance, and 5 error limit for all scan filters. The *m/z* values of the targeted precursor ions are shown on top of the respective SSN. For experimental details see Dataset 1 in Supplementary Table 9.

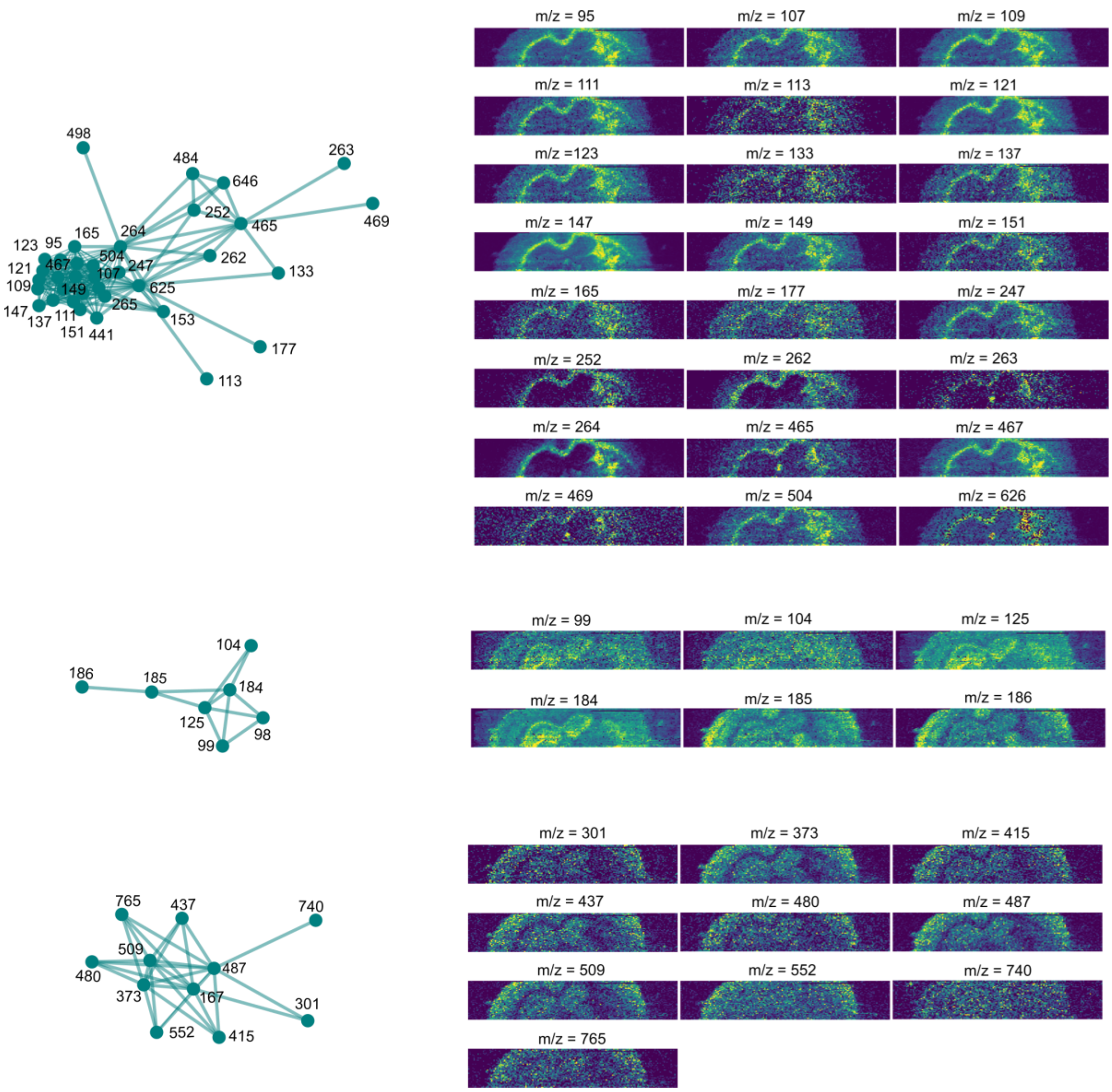

**Supplementary Figure 6. Product ion network clusters and corresponding images from fragmentation of the isolation window centered at 810.60 ± 0.35 Da.** The product ion images were acquired using the PIA of a mouse brain tissue section. The subsequent SSN reveals that product ions are separated into three groups based on their spatial distribution, as shown by the orange and blue SSN, suggesting the presence of at least three structurally different precursors. For experimental details see Dataset 1 in Supplementary Table 9.

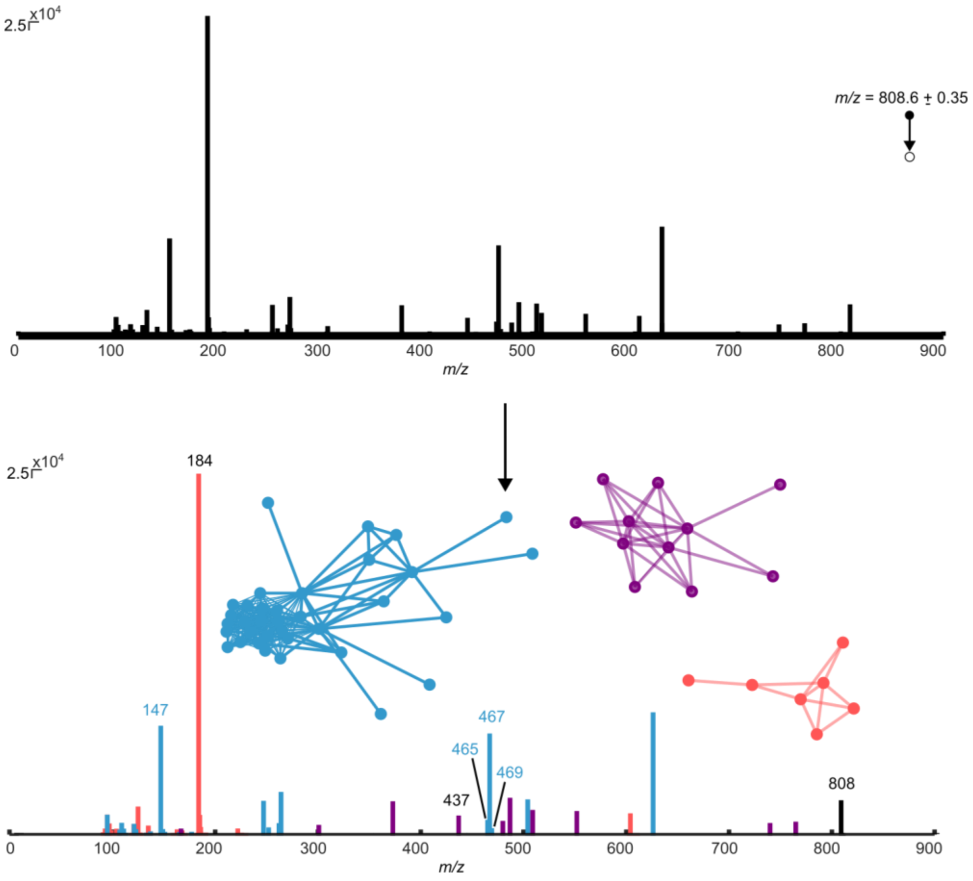

**Supplementary Figure 7. SSN works as a separation dimension unique to MSI and deconvolutes product ion mass spectra.** The convoluted ITMS$^2$ mass spectrum produced from the product ions of all precursors in the 808.6 ± 0.35 window is readily deconvoluted by SSN. The product ions network clusters shown in blue, purple, and red correspond to the blue, purple, and red product ions in the deconvoluted mass spectrum, respectively. For experimental details see Dataset 1 in Supplementary Table 9.

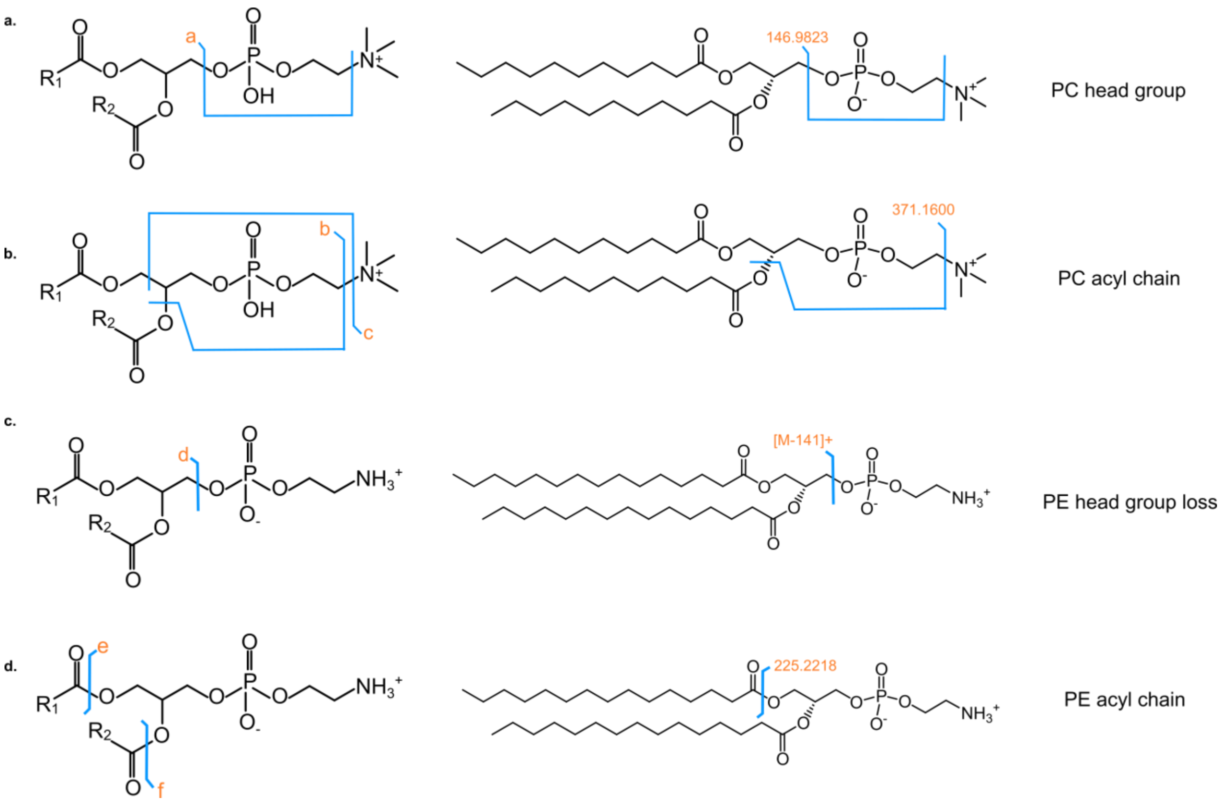

**Supplementary Figure 8. Fragmentation patterns of phosphotidylcholine (PCs) and phosphotidylethanolamine (PEs) as sodiated adducts in HCD.** The PC head group is detected at m/z 146.9823 after to loss of the acyl chains and the choline group (a). After the loss of choline and one acyl chain, either the sn-1 or sn-2, the *m/z* of the remaining head group, the glycerol backbone and the remaining acyl chain is detected (b). The neutral loss of the PE head group (-141) results in the detection of the acyl chains and the glycerol backbone (c). Either the sn-1 and sn-2 acyl chains of the PE are detected after the fragmentation in the ester linkage (d). Note that both PC and PE fragmentation in HCD produce diagnostic product ions corresponding to the acyl sidechain as well as information on head group loss in positive ion mode. Contrarily, collision induced dissociation (CID) only provides information on head group loss.

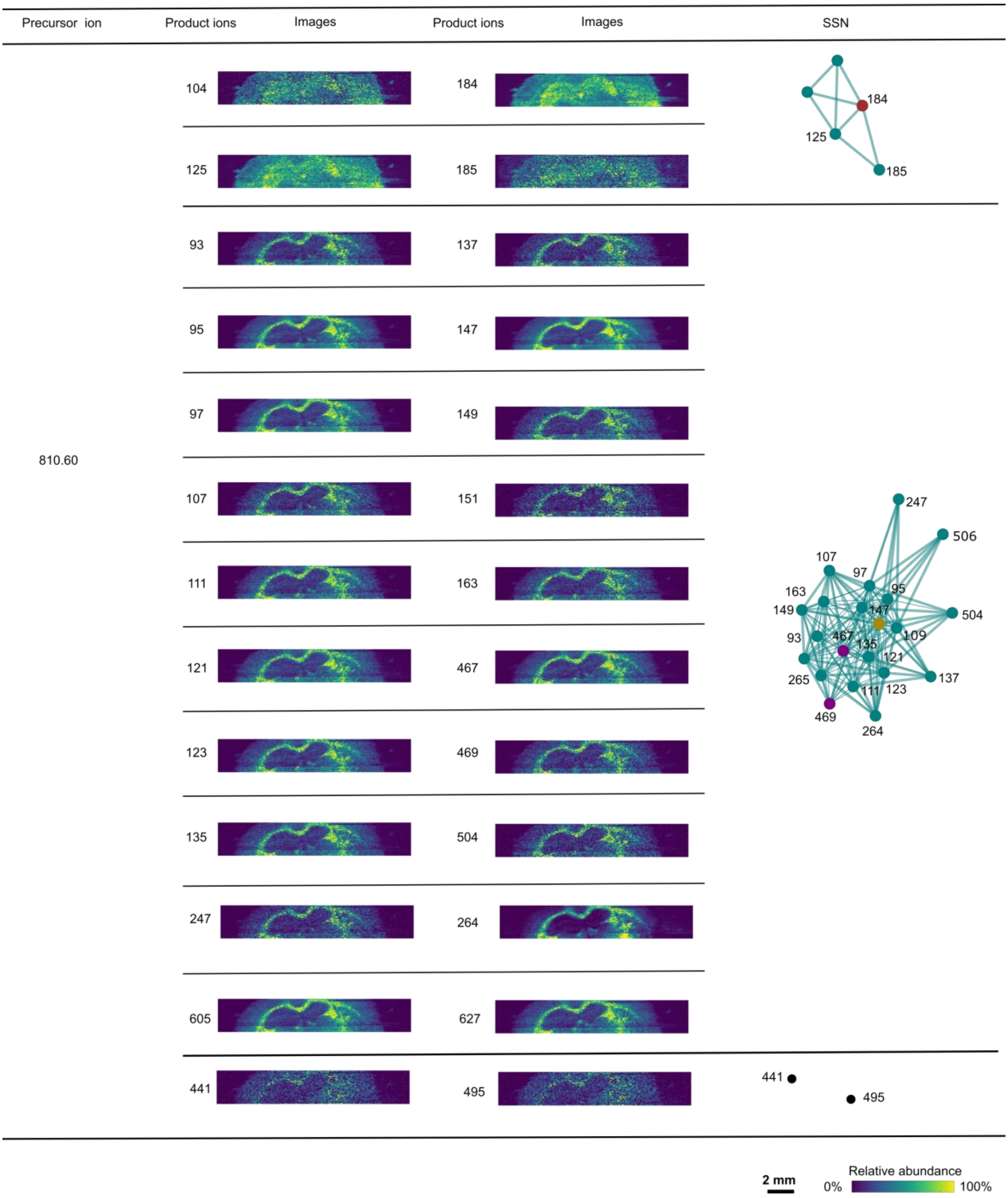

**Supplementary Figure 9. Product ion image networks from PIA SSN.** The *m/z* values and corresponding product ion images were acquired from the precursor window of 810.60 ± 0.35 Da using the PIA of a mouse brain tissue section. The subsequent SSN reveals that product ions are separated into two groups based on their spatial distribution, as shown by the orange and blue SSN, suggesting the presence of two structurally different precursors. Two additional disconnected nodes are present, indicating the presence of a third, but less abundant, precursor. From top to bottom, the represented ions are [PC 38:4+H]$^+$, [PC 18:0_18:1+Na]$^+$, and [PC 16:0_20:1+Na]$^+$. For experimental details see Dataset 1 in Supplementary Table 9.

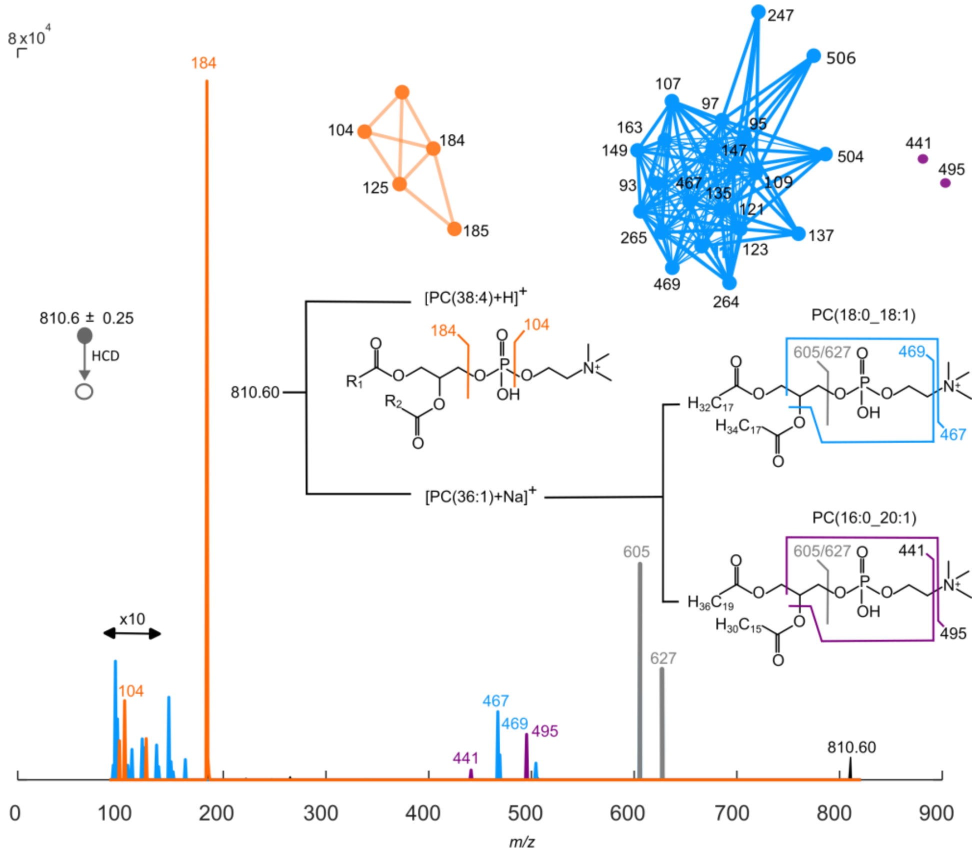

**Supplementary Figure 10. SSN deconvolutes product ion mass spectra and facilitates annotation of isobars and isomers.** The convoluted ITMS$^2$ mass spectrum produced from the product ions of all precursors in the 810.6 ± 0.35 window is readily deconvoluted by SSN. The product ions corresponding to the blue and orange SSN are shown in blue and orange, respectively. This shows that the two distinct isobars of $[PC(38:4)+H]^+$ and $[PC(36:1)+Na]^+$, which have different spatial distributions, both originate from the precursor ion at *m/z* 810.60. Furthermore, the data reveals that $[PC(36:1)+Na]^+$ consists of the two isomers PC(18:0_18:1) and PC(16:0_20:1). For experimental details see Dataset 1 in Supplementary Table 9.

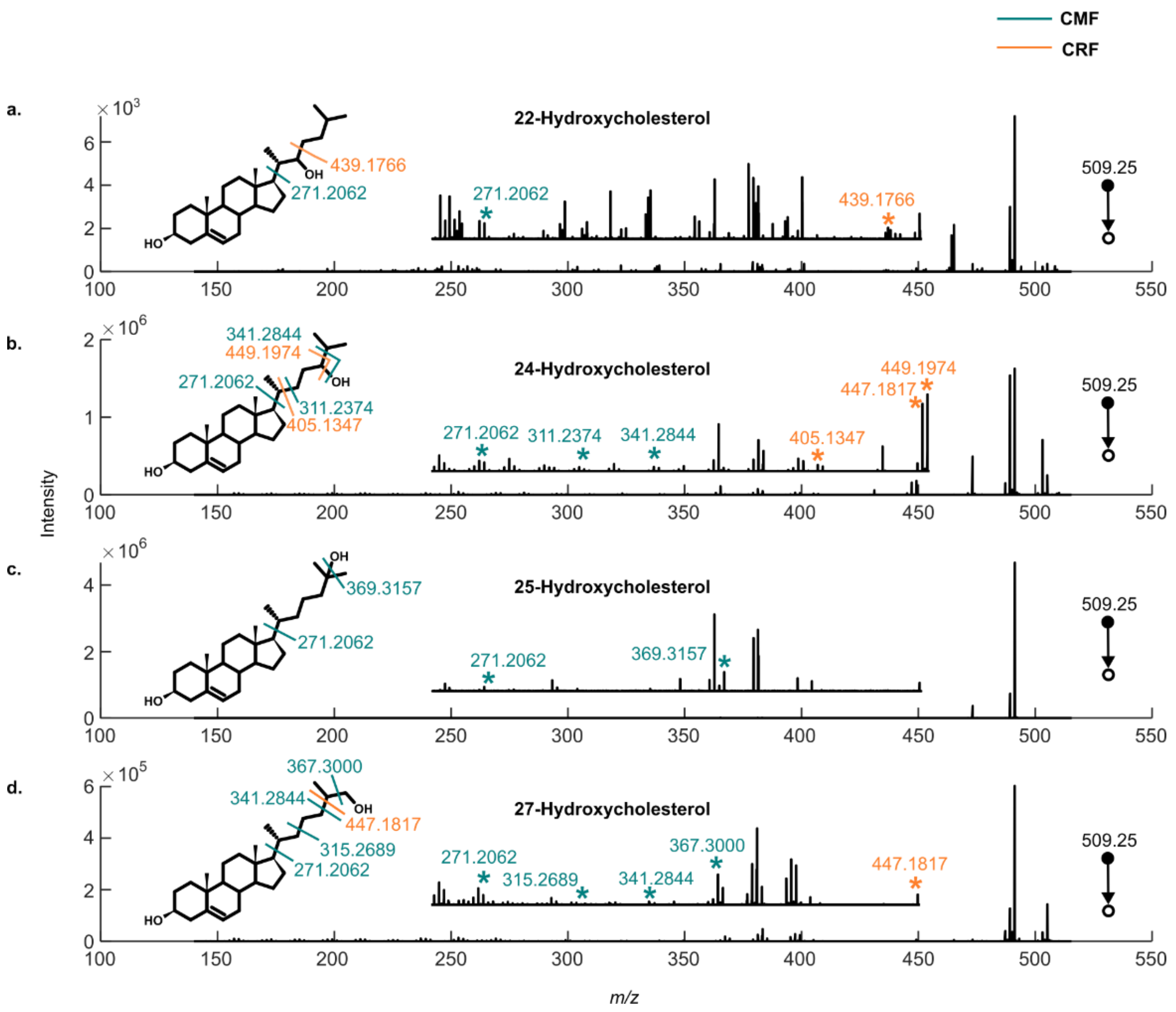

**Supplementary Figure 11. Fragmentation of argenated adducts of hydroxycholesterols**. Consistent fragmentation sites are identified for argenated hydroxycholesterols upon HCD. $FTMS^2$ mass spectrum of 5 µM standards of (a) 22-hydroxycholesterol, (b) 24-hydroxycholesterol, (c) 25-hydroxycholesterol, and (d) 27-hydroxycholesterol. The annotated product ions are generated by either charge migration fragmentation (CMF, in blue) or charge retention fragmentation (CRF, in orange). In CRF $Ag^+$ is retained on the product ions while in CMF $Ag^+$ is lost during fragmentation.

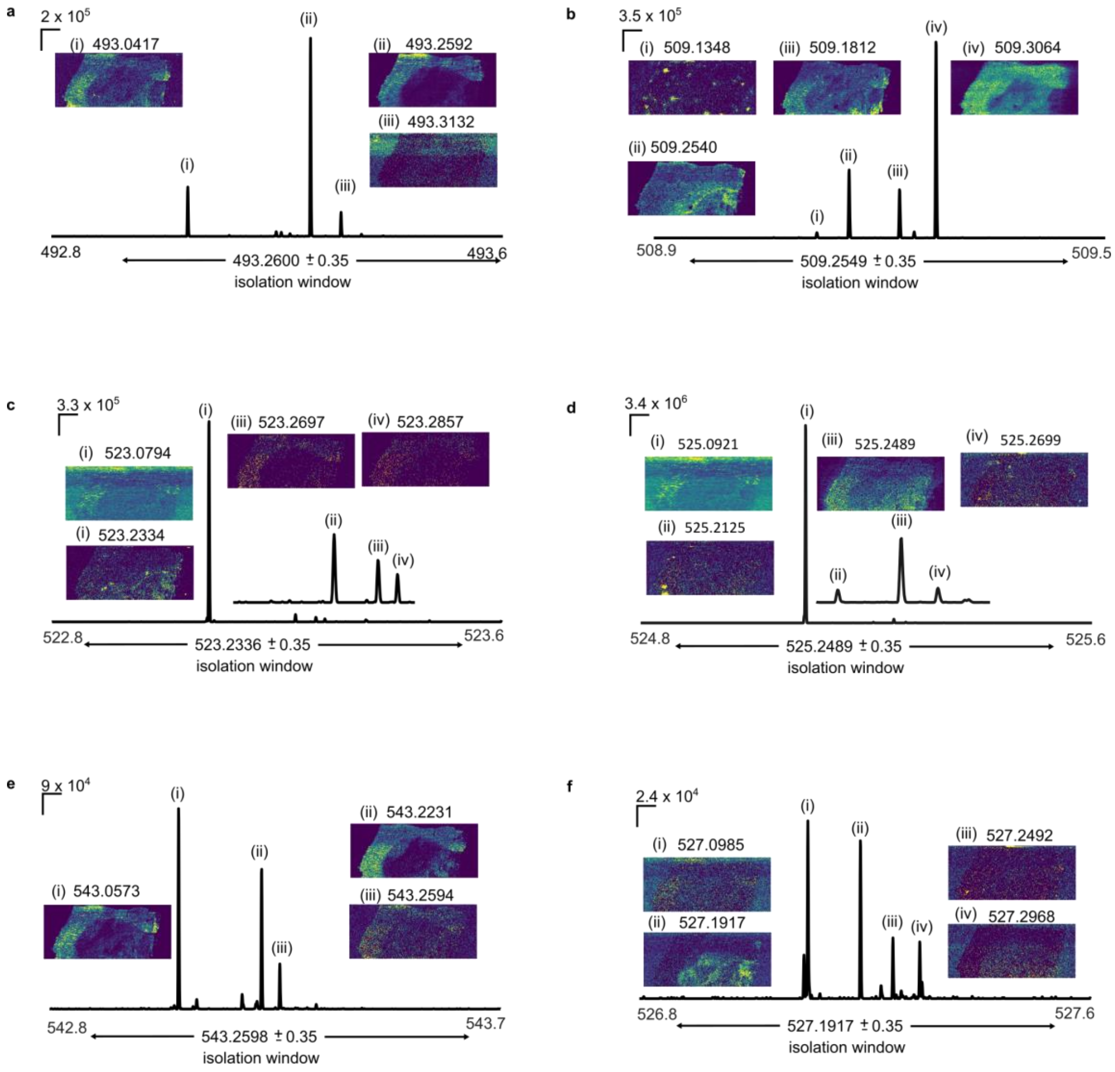

**Supplementary Figure 12. The isolation window contains multiple precursors ions**. FTMS spectra detailing the isolation window of precursor ions selected for $ITMS^2$. The isolation windows for cholesterol and oxidized cholesterol products (ST1-4 and ST6-7) are displayed in a-f. Data is acquired from in human multiple sclerosis brain tissues and show the high complexity of detected ions in the selected regions. The ion images show the different distributions that arise from the respective precursors (i-iv) in the isolation windows. The co-isolation of multiple precursors increases complexity when decoding $ITMS^2$ spectra**.** For experimental details see Dataset 6 in Supplementary Table 9.

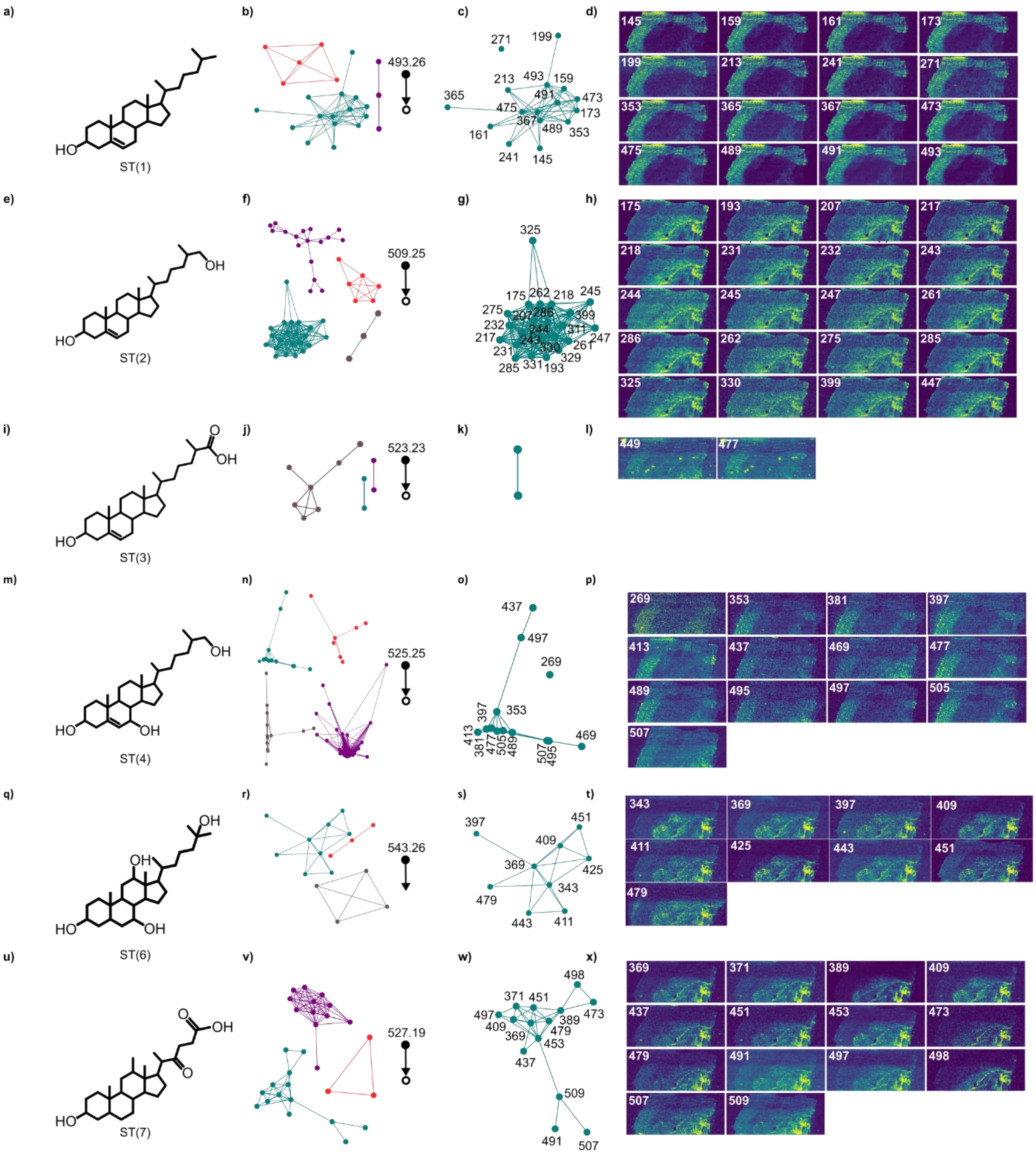

**Supplementary Figure 13. The SSN deconvolutes complex ITMS$^2$ spectra produced by the fragmentation of co-isolated precursor ions.** Characterization of oxysterol structures detected by PA nano-DESI MS$^2$I of human multiple sclerosis brain tissue. Structural annotation of cholesterol and oxidized cholesterol products using HCD of $[M+^{107}Ag]^+$ adduct in ITMS$^2$. a-d cholesterol, e-h hydroxycholesterol i-l hydroxycholestenoic acid, m-p dihydroxycholesterol, q-t cholestantetraol, u-x homodeoxycholic acid. For each targeted molecule, SSN and all product ions in the SSN are provided. One additional structural annotation (dihydroxy-cholestenoic acid) is shown in figure 2. For experimental details see Dataset 5 in Supplementary Table 9.

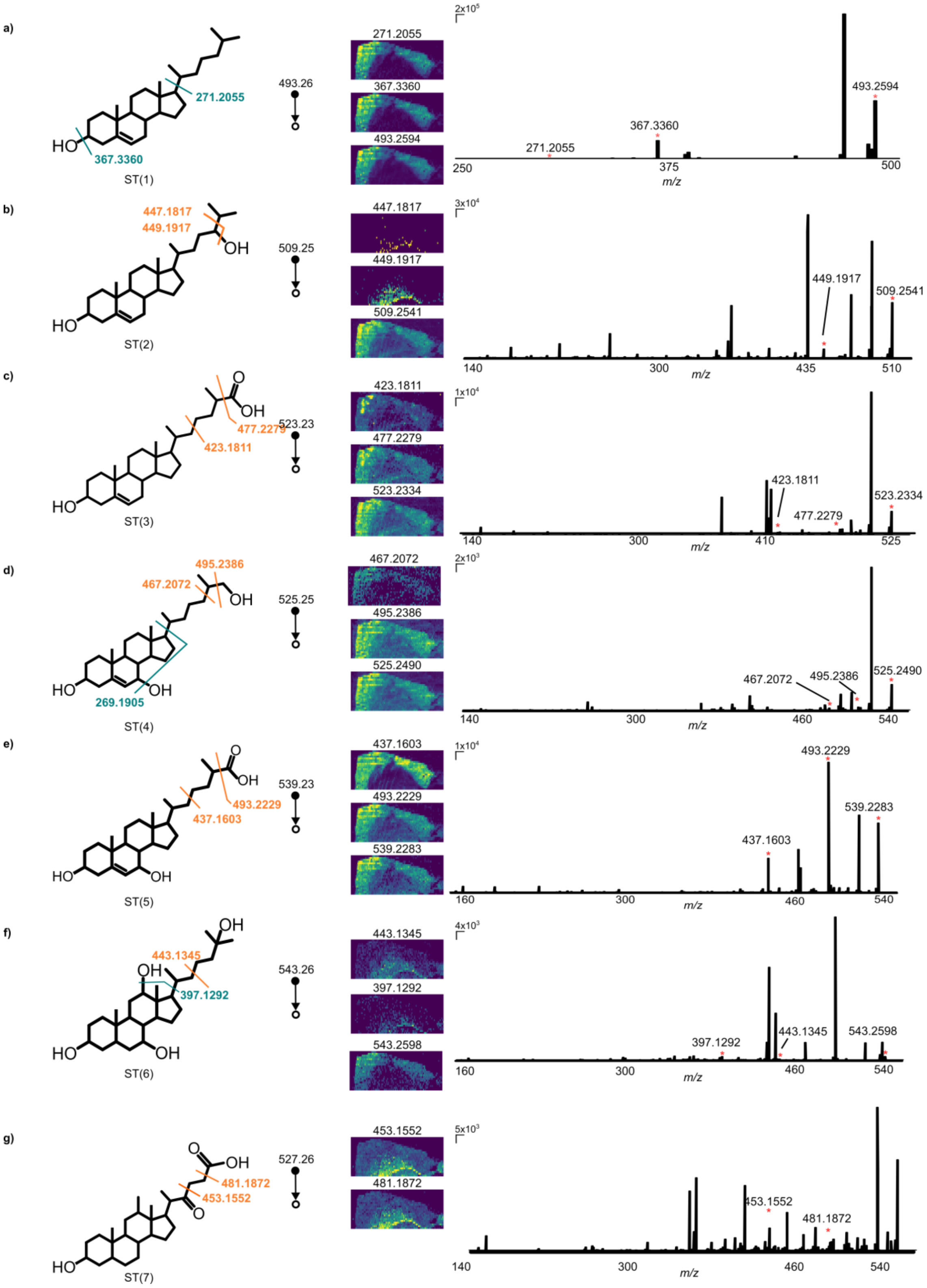

**Supplementary Figure 14. Detected oxysterol species from human multiple sclerosis tissue by sequential FTMS²I.** High mass resolution imaging of detected diagnostic product ions from cholesterol and cholesterol oxidation products (ST1-7) together with the respective fragmentation sites, precursor and product ion images, and FTMS² spectra are given in a-g. The details show that the mass channel of ST1, ST3, and ST4 have identical ion images for the product ions and the precursor ion (a, c-d) and the mass channel of ST2, ST5 and ST6 (b, e-f) may have multiple isomers. The abundance of the precursor ion for ST7 (g) is under detection limit. All co-isolated precursor ions show different distributions (Supplementary Fig. 11). For experimental details see Dataset 6 in Supplementary Table 9.

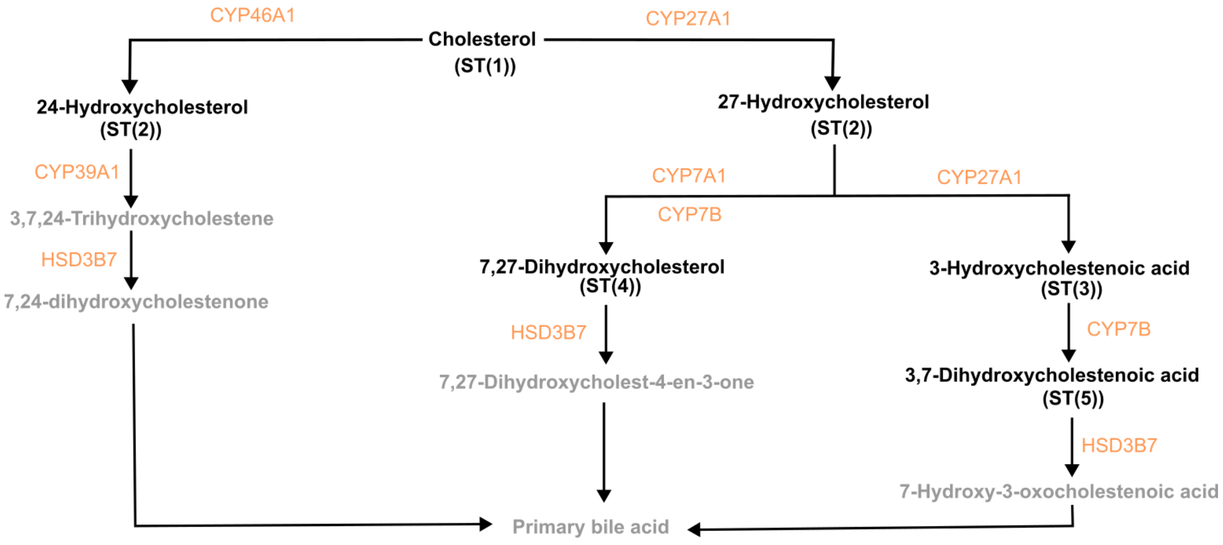

**Supplementary Figure 15. Metabolic pathway for biosynthesis of bio acids from cholesterol**. The molecules highlighted in black with the abbreviations are detected in the human brain tissue, while the grey species are not detected. The responsible enzyme for each biochemical conversion is detailed in orange. The schematic is made from the KEGG database.

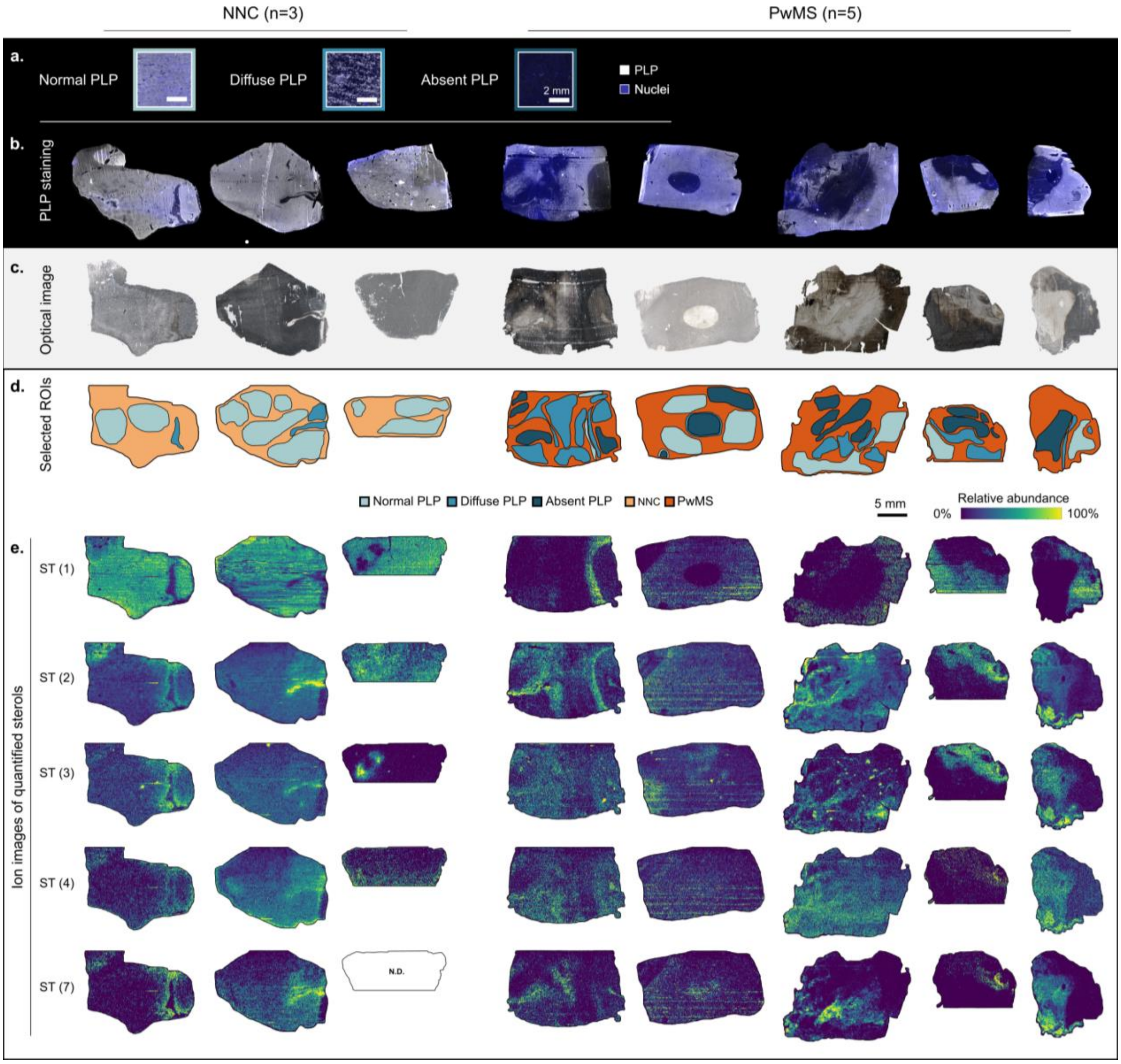

**Supplementary Figure 16. Multimodal imaging of cholesterol and oxidized cholesterol products in human white matter brain tissue sections.** Representative classifier images of fluorescent proteolipid protein (PLP) immunoreactivity were acquired as stated in the Methods section. Scalebar is 2 mm. (a) Full tissue scans of PLP immunoreactivity of human brain tissues (NNC=3, PwMS=5). Image intensities are scaled individually. (b) Brightfield microscopy images acquired using a Meyer Instruments PathScan Enabler IV slide scanner device. (c) Resulting ROIs for each tissue section, colored according to the PLP classification (light blue: normal PLP, medium blue: diffuse PLP, dark blue: absent PLP). (d) Ion images of cholesterol and oxidized cholesterol products in human white matter brain tissues from NNC and PwMS normalized to TIC and the 99$^{th}$ percentile intensity. Scalebar is 5 mm and applies to images b-e. N.D.: not detected. Subsequent tissue sections are used for PLP and MSI imaging modalities.

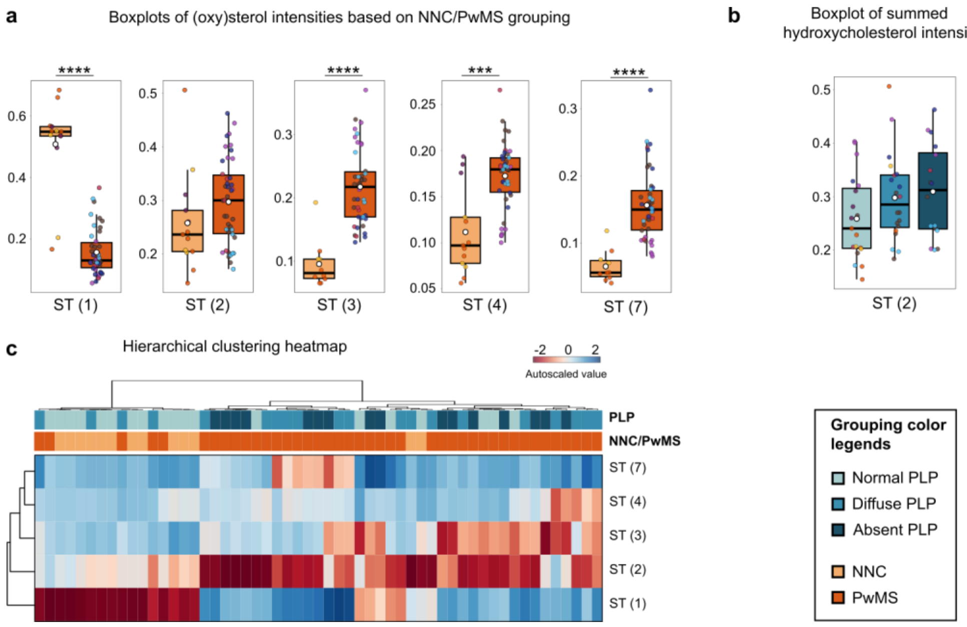

**Supplementary Figure 17. Differentiation and clustering of brain tissue regions based on their oxysterol profiles. (a)** Boxplots representing the fractional abundance of detected sterols between ROIs corresponding to NNC (light blue) and PwMS (orange) subject samples. Note that since quantitation is based on $MS^1$, the two detected isomers 24-hydroxycholesterol and 27-hydroxycholesterol are denoted together as ST(2). All other presented mass channels are confirmed by $FTMS^2I$ to only contain the annotated sterol. Significant differences were observed for all the molecules except ST(2). To define significance, Wilcoxon test with Benjamini-Hochberg FDR correction was performed (*:p<0.05, **:p<0.01, ***:p<0.001, ****:p<0.0001). Coloring of points displays data from individual subjects as denoted by Supplementary Table 8. **(b)** Boxplot representing the fractional abundance of ST(2) showing non-significant differences between ROIs corresponding to normal PLP (light blue), diffuse PLP (medium blue), and absent PLP (dark blue) subject samples. For additional boxplots, see Figure 5 in the main text. **(c)** Hierarchical clustering heatmap of (oxy)sterols across all ROIs shows a slight differentiation of normal PLP ROIs from diffuse and absent PLP ROIs and a considerable differentiation when the grouping is based on the subjects (NNC vs PwMS). For clustering, Ward method with Euclidian distance measures was used and values were autoscaled within each ROI sample. The coloring of hues for the autoscaled values is displayed in the top right corner.
